# Aging alters the distribution, stability, and transcriptional signature of engram cells

**DOI:** 10.1101/2025.08.29.673010

**Authors:** Miguel Fuentes-Ramos, Marta Alaiz-Noya, Federico Miozzo, Alba Vieites-Prado, Jordi Fernández-Albert, Nicolas Renier, Angel Barco

## Abstract

While aging impairs memory precision, its effects on engram dynamics and gene expression remain poorly understood. To address this, we used TRAP2 activity-reporter mice, nuclear tagging, and FOS-based activity mapping to track neurons activated during contextual fear memory encoding and reactivated during recall in young and aged mice. Across 378 brain regions, we quantified engram size, spatial distribution, and reactivation stability. We further applied fluorescence-activated nuclear sorting (FANS) combined with single-nucleus RNA sequencing (snRNA-seq) to characterize gene expression changes associated with memory encoding and recall across diverse cell types. In addition, we compared the transcriptional profiles of first-time versus second-time neuronal responder cells in the dentate gyrus. Aged brains exhibited altered engram allocation, reduced reactivation stability, and distinct gene expression patterns during memory retrieval. These findings reveal age-related changes in the organization and molecular identity of memory traces, providing mechanistic insight into cognitive decline and highlighting potential targets for intervention.

## Introduction

Memories are deposited in discrete neuronal ensembles in the hippocampus, amygdala, and other brain regions involved in memory formation(*1*, *2*). The cells in those ensembles are known as engram cells and are thought to undergo enduring physical or chemical changes initiated during learning that enable their selective reactivation during retrieval to produce the recall of the experience(*3*). Despite progress in memory trace transcriptomics, the expression changes in engram cells linked to memory persistence or age-related cognitive decline are still unclear(*4*).

The Targeted Recombination in Active Populations (TRAP) mouse strains enable high temporal resolution and permanent tagging of neurons activated during a specific experience, facilitating longitudinal analyses of activity-induced changes even long afterward(*5*). These strains utilize a tamoxifen (TAM)-dependent CreERT2 recombinase driven by the activity-dependent promoters of immediate early genes (IEGs) such as *Arc* or *Fos*(*6*). An improved version of the TRAP system, known as FosTRAP2(*7*, *8*), enhanced sensitivity for activity labelling compared to the FosTRAP(1) mouse line and reduce leakiness observed in the ArcTRAP mouse line(*8*).

In this study, we used TRAP2 mice to investigate the distribution, cellular composition, and reactivation stability of memory engrams across 378 brain regions in both young and aged mice. Our analysis focused not only on neurons labeled during memory formation but specifically on those reactivated during retrieval, which remains poorly characterized(*9*). To achieve this, TRAP2 mice were crossed with a Cre recombinase-dependent reporter, SUN1-GFP, which labels the nuclear envelope, facilitating automated quantification and isolation of the activated cells. Using immunolabeling-enabled three-dimensional imaging of solvent-cleared organs (iDISCO(*10*)) and ClearMap2(*11*, *12*) software, we analyzed and mapped engram cells throughout the main brain regions involved in memory. Additionally, we combined TRAP2 mice with fluorescence activated nuclear sorting (FANS) and single nucleus RNA-seq (snRNA-seq) to investigate transcriptional signatures induced by memory retrieval. Our analyses identified granule neurons in the dentate gyrus (DG) as the most affected population, prompting a targeted nuclear RNA-seq (nuRNA-seq) screen comparing neurons activated exclusively during encoding, recall, or both. This work revealed significant differences in reactivated cells and uncovered age-related changes in engram activation between young and aged mice, providing new insights into the molecular mechanisms underlying age-associated cognitive decline.

## Results

### A fluorescence reporter system for efficient brain-wide engram mapping

FosTRAP2 mice offer a sensitive and efficient strategy to label active neurons(*6–8*) (**Sup. Figure S1A**). However, TRAP2-mediated tagging occurs at lower efficiency than endogenous *Fos* expression(*6*). To develop a more robust and versatile method, we introduced two recombination-dependent reporters, SUN1-GFP and tdTomato, both knocked into the Rosa26 locus on different alleles within the same animals. Initial validation in mice constitutively expressing CreERT2 in principal neurons established the optimal time window (**Sup. Figure S1B-D**) and confirmed that a single dose of 4-OHT was sufficient to induce recombination in both alleles (**Sup. Figure S1E-G**). In contrast, when combined with TRAP2 mice (**Figure 1A**), the double-reporter system revealed that most cells labeled following contextual fear conditioning (CFC) expressed only one reporter (**Figure 1B-D**). A similar pattern was observed following kainic acid (KA)-induced seizures, although this paradigm resulted in a higher number of neurons labeled with either or both reporters (**Figure 1E)**. Probabilistic analysis indicated that inefficient stochastic recombination, rather than allelic exclusion, accounts for the sparse dual labeling (**Sup. Figure S1H**). Given that tagging efficiency appeared constrained by Cre recombinase levels and that the *Fos* promoter is weaker than CamK2a, we generated mice homozygous for both *Fos*TRAP2 and SUN1-GFP alleles (hereafter referred to as NucTRAP2). Homozygosity markedly increased labeling efficiency with a single dose of 4-OHT (**Figure 1F-G**) without affecting behavior (**Sup. Figure S1I**). Notably, NucTRAP2 mice exhibited a significantly higher number of labeled neurons compared to heterozygotes (**Figure 1H**), reaching levels comparable to endogenous FOS expression (**Figure 1I**). To assess baseline recombination in the absence of 4-OHT, we examined reporter expression during novel environment exploration at 2 and 16 months of age. Spontaneous recombination remained negligible even in aged animals(*6*, *13*) (**Supp. Figure S1J**). Altogether, NucTRAP2 mice significantly improved current engram tagging systems, closely matching endogenous Fos activation.

**Figure 1.**
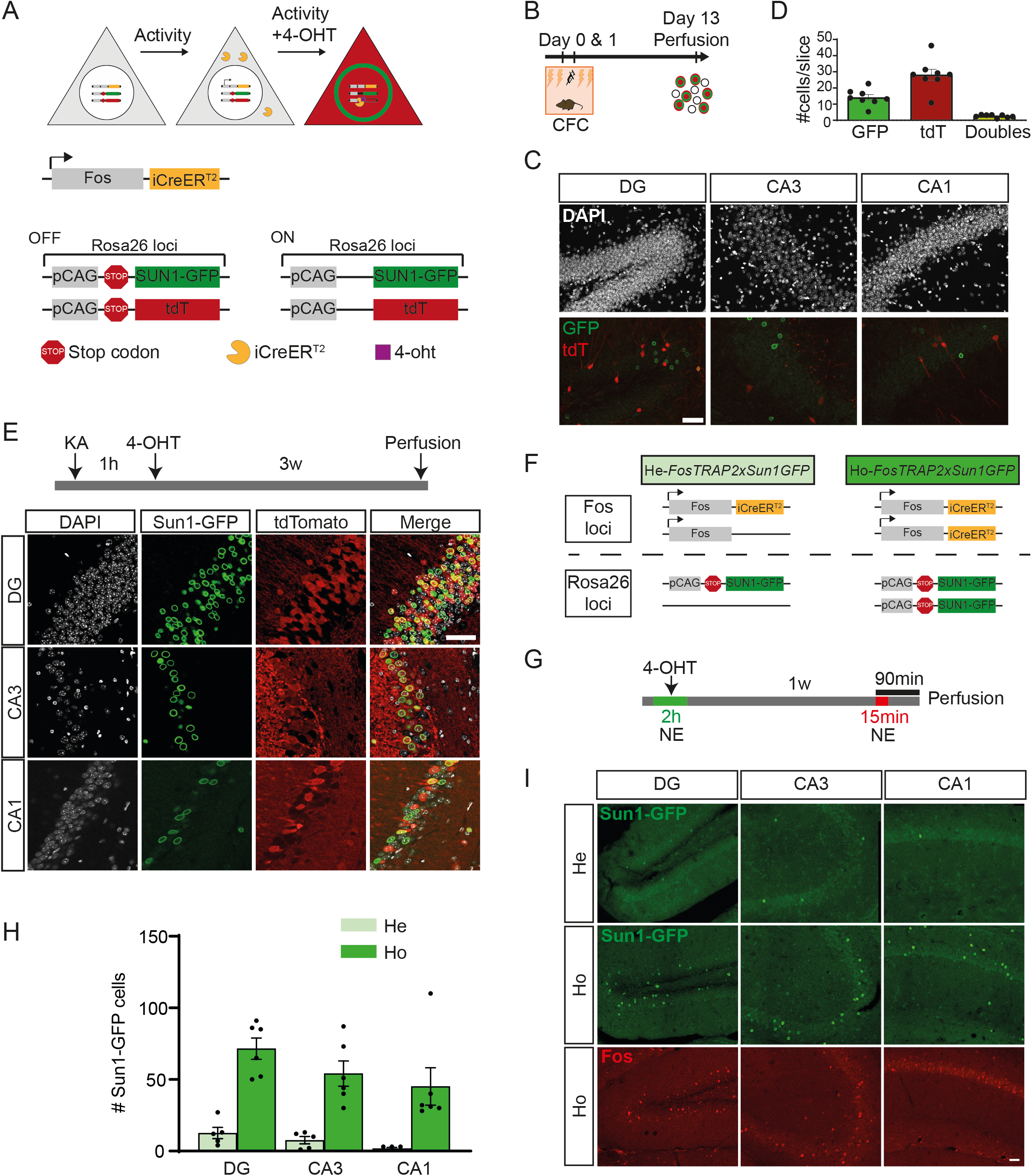
TRAP2 characterization and optimization of activity-dependent genetic tagging. **A.** FosTRAP2 animals were crossed with mice bearing a SUN1-GFP reporter transgene knocked into one allele of Rosa26 locus and a tdTomato reporter transgene in the other to produce double knock-in mice. Both reporters are Cre-recombination-dependent, enabling visualization of the nuclear envelope (SUN1-GFP) or the entire neuronal morphology (tdTomato) upon *Fos* activation and 4-OHT administration. **B-D.** *FosTRAP2xSUN1GFP* mice underwent fear training over two consecutive days. **B.** The schematic outlines the experimental design for CFC-induced neuronal activation. **C.** Representative confocal images of the hippocampus. **D.** Quantification of cells expressing each reporter. **E.** Top: Experimental design for KA-induced neuronal activation. Bottom: Representative confocal images from *FosTRAP2xSUN1GFP* mice one week after KA administration. **F.** Comparison of *FosTRAP2xSUN1GFP* homozygous and heterozygous. **G.** Experimental design to evaluate the levels of reporter labelling between *FosTRAP2xSUN1GFP* homozygous and heterozygous. **H.** Quantification of GFP-expressing neurons in *FosTRAP2xSUN1GFP* homozygous and heterozygous mice. **I.** Representative confocal images comparing SUN1-GFP tagging in *FosTRAP2xSUN1GFP* homozygous and heterozygous mice with endogenous *Fos* induction. Scale bars 50 µm.

### Brain-wide mapping of activated cells during memory encoding and recall

While *Fos* expression during memory encoding and retrieval has been studied in specific brain regions, its brain-wide distribution remains unclear. To address this, we used NucTRAP2 mice, in which strong nuclear envelope labeling facilitates automated quantification of tagged neurons. Mice were subjected to an extended CFC paradigm (120 min), with 4-OHT administered midway through training. As a control, littermates received 4-OHT without undergoing training and remained in their home cages (HC). Seven days later, brains were cleared using iDISCO and imaged for SUN1-GFP expression with ClearMap2 (**Figure 2A**). Quantification across 378 brain regions (**Supp. Table 1**) revealed widespread activation after conditioning, with pronounced differences between CFC and HC groups (**Figure 2B**; **Supp. Figure S2A-B**). For instance, CFC induced robust labeling across the isocortex and multiple subcortical structures (**Supp. Figure S2C**, **Supp. Table 1**). Correlation analysis across brain regions further indicated that CFC triggered coordinated activation particularly within the hippocampal formation and cortical subplate (**Figure 2C**), suggesting synchronized engagement of distributed memory-related circuits during associative learning.

**Figure 2.**
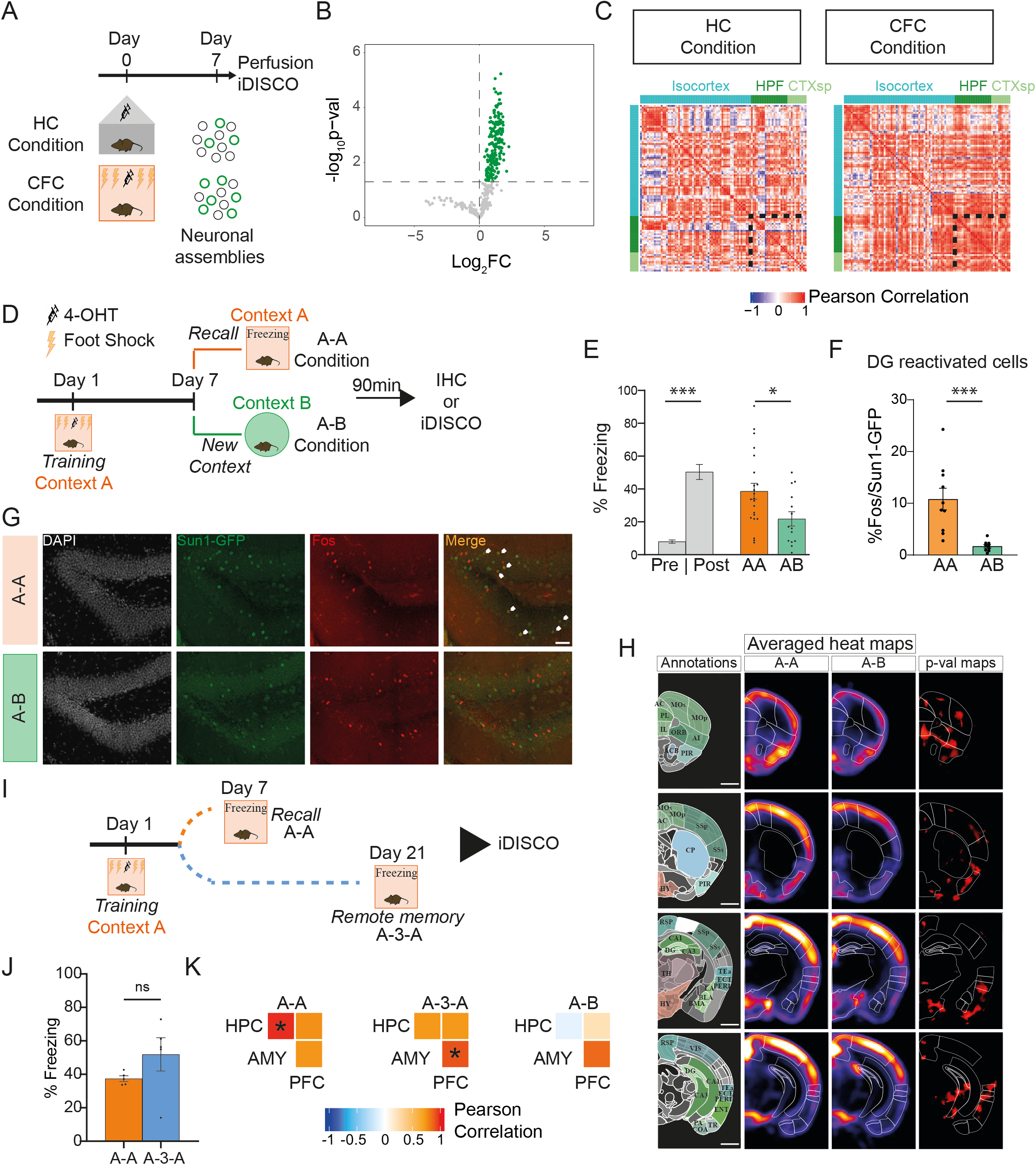
Tagging of a CFC engram. **A.** Experimental design to study activity-dependent labelling system. 4-OHT was administrated during a contextual fear training (CFC, n=5) or in home cage (HC, n=5) to label neuronal activity in each moment. Seven days after brains were collected and whole-mount iDISCO+ immunolabeling was performed. **B.** Volcano plot showing the CFC-induced regions as log2 fold change in SUN1-GFP+ cells (CFC vs HC group; p-val < 0.05). **C.** A subregion-by-subregion matrix of brain activity is obtained by pairwise correlation of the regional patterns of SUN1-GFP expression for each experimental group. **D.** Experimental design to study specificity of the contextual fear memory. Seven days after contextual fear conditioning in Context A mice were exposed to same (A-A) or different context (A-B) to evaluate behavioural responses and cellular activation in both moments. **E.** Freezing behaviour during memory recall (A-A: n=22; A-B: n=14; Mann–Whitney U Statistic). **F-G.** Quantification (F; A-A: n=10; A-B: n=9; Mann–Whitney U Statistic) and representative images of reactivated cells (SUN1-GFP^+^ and Fos^+^) in dentate gyrus for A-A and A-B. Scale bar, 50 µm. **H.** Coronal views of the automated analysis of Fos^+^ cell distribution. Panels show the reference annotations, averaged density maps for each condition (5 brains averaged per condition) and p-value maps (red: increased Fos expression in the A-A group; green: increased Fos expression in the A-B group). Scale bar, 500 µm. **I.** Experimental design to study the stability of the engrams over time. Mice were re-exposed to Context A one (A-A) or three weeks (A-3w-A) after CFC. **J.** Freezing behaviour during memory recall (A-A: n=5; A-3w-A: n=5; Mann–Whitney U Statistic). **K.** Correlation between Fos levels of the three most studied regions involved in contextual memory: hippocampus (HPC: granular-polymorph-molecular layers of DG, CA1, CA2 and CA3), amygdala (AMY: basolateral, basomedial and lateral amygdala) and prefrontal cortex (PFC: anterior cingulate, prelimbic, infralimbic and orbital areas). \**P* < 0.05, \*\**P* < 0.01, \*\*\**P* < 0.001, ns = no significant.

Next, we compared cell activation during training and retrieval in two experimental groups. Seven days after TRAPing engram cells during CFC, one group was re-exposed to the same context (A–A group), while the other was exposed to a novel context (context B; A–B group) (**Figure 2D**). As expected, mice re-exposed to the conditioned context exhibited significantly higher freezing (**Figure 2E**, **Supp. Figure S3A**) and tagged cells reactivation at the DG (**Figure 2F-G**), a region critical for contextual discrimination(*14*), consistent with context-specific tagging. Notably, the overall number of FOS+ neurons did not differ between groups (**Supp. Figure S3B**). Expanding the analysis brain-wide, also revealed similar overall levels of activation in the A–A and A–B groups (**Supp. Figure S3C**). However, recall of the conditioned context engaged a broader network, with 38 additional regions activated in the A–A group (p < 0.05; **Figure 2H**, **Supp. Figure S3D-E** and **Supp. Table 2**). These included canonical memory regions such as the basomedial amygdala, entorhinal cortex, anterior cingulate cortex, and infralimbic cortex (**Supp. Figure S4**). Intriguingly, several regions not traditionally associated with mnemonic processes, including the agranular insular cortex, ventral posteromedial thalamic nucleus, and supramammillary nucleus, also showed enhanced activation during recall (**Supp. Figure S4**). Together, these findings suggest that memory retrieval engages a more distributed and expansive brain network than previously appreciated, extending beyond classical memory circuits.

### Stability of engram cells

To assess the temporal stability of engram cells, we compared neuronal reactivation in mice re-exposed to the conditioned context either 1 or 3 weeks after fear conditioning (**Figure 2I**). Freezing behavior during recall was comparable between groups (**Figure 2J, Supp. Figure S5A**), and FOS expression patterns were similarly stable (**Supp. Figure S5B-C**), with only minor regional differences detected (**Supp. Figure S5D** and **Supp. Table 3**). These results suggest that recall-induced activation patterns remain largely stable over time. However, comparisons of 1- and 3-week recall with novelty exploration revealed less than 25% overlap (**Supp. Figure S5E-F**), suggesting that dynamic changes accompany memory consolidation. This is consistent with models proposing progressive reorganization of memory traces across brain networks(*15*, *16*), wherein hippocampus-dependent memories gradually shift toward increased cortical engagement. In fact, whole-brain analyses revealed that during 1-week recall, FOS ensembles exhibited stronger coupling between the hippocampus and amygdala, while by 3 weeks, the strongest correlations emerged between the amygdala and prefrontal cortex (**Figure 2K**; **Supp. Figure S5G**). No comparable ensemble coupling was observed following exploration of a novel context (**Figure 2K**; **Supp. Figure S5G**).

### Age-related memory deficits are associated with impaired network activation

To investigate the impact of aging on engram mapping and stability, we subjected aged (18-22 months) NucTRAP2 mice to the extended CFC protocol (**Figure 3A**). No significant differences in distance travelled during novel environment exploration (**Supp. Figure S6A**) or during fear training (**Supp. Figure S6B**) were observed between young and aged mice, although minor differences in delta freezing suggested that young mice exhibited a stronger response during conditioning (**Supp. Figure S6C**). During retrieval, aged mice failed to preferentially freeze in the conditioned context, indicating impaired context discrimination (**Figure 3B**; **Supp. Figure S6C-D**). Classical CFC confirmed these findings (**Supp. Figure S6E–H**). Additionally, aged mice showed greater behavioral variability, with a markedly higher coefficient of variation in the Discrimination Index compared to young mice (Day 7: young CV = 54.96%, aged CV = 204.7%; Day 8: young CV = 101.3%, aged CV = 141.0%), consistent with individual variability in age-related cognitive decline reported previously(*17*, *18*).

**Figure 3.**
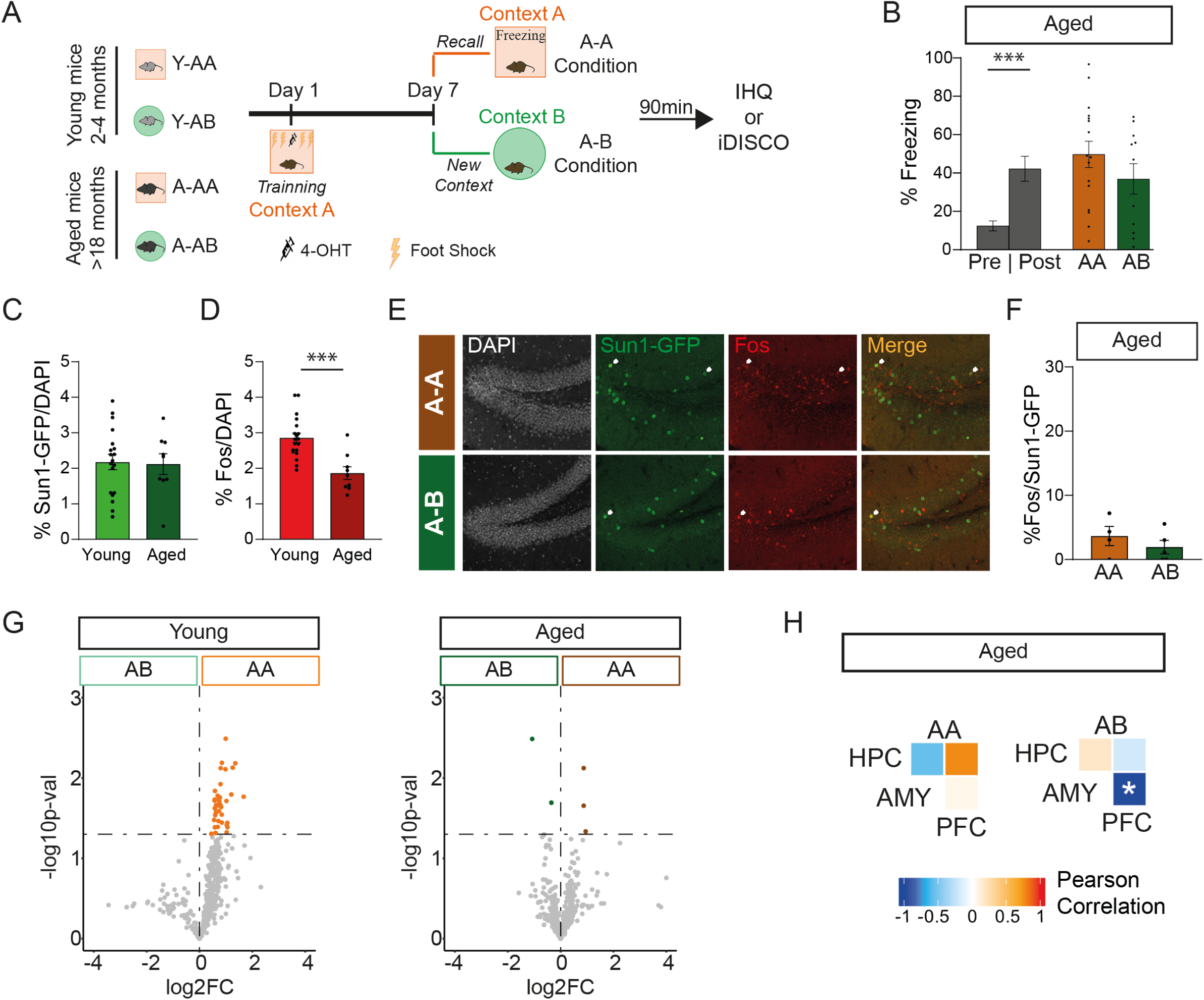
Age-associated imprecise memory and different patterns of engram activation in aged mice. **A.** Experimental procedure. **B.** Freezing behaviour during memory recall (A-A N=16; A-B N=11; Mann–Whitney U Statistic). **C-D.** Quantification of the percentage of SUN1-GFP+ (C; young N=19; aged N=9; Mann–Whitney U Statistic) and Fos+ (D; young N=19; aged N=9; Mann–Whitney U Statistic) cells relative to the total number of DAPI+ cells in DG of young and aged mice. **E-F.** Representative images and quantification (F; A-A N=4; A-B N=5; Mann–Whitney U Statistic) of reactivated cells (SUN1- GFP + & Fos+) in dentate gyrus for A-A and A-B in aged mice. **G.** Volcano plot of fear recall-induced regions in young (left) and aged (right) mice as fold change in Fos+ cells (A-A over A-B). Significant regions appearing coloured. **H.** Correlation between Fos levels of the three most studied regions involved in contextual memory in aged mice: hippocampus (HPC: granular-polymorph-molecular layers of Dentate Gyrus, CA1, CA2 and CA3), amygdala (AMY: basolateral, basomedial and lateral amygdala) and prefrontal cortex (PFC: anterior cingulate, prelimbic, infralimbic and orbital areas). \**P*<0.05, \*\**P*<0.01, \*\*\**P*<0.001.

To determine whether behavioral deficits correlated with altered engram mapping, we compared the number of SUN1-GFP+ engram cells in the DG of young and aged NucTRAP2 mice following CFC. Although the total number of tagged cells did not differ between groups (**Figure 3C**), aged mice exhibited impaired activation and reactivation of DG engram cells during context-specific recall (**Figures 3D-F**), indicating reduced context specificity at the neuronal level. Furthermore, brain-wide FOS mapping revealed that aged mice displayed greater similarity in *Fos* expression patterns between conditioned and novel contexts relative to young mice (**Figure 3G** and **Supp. Table 4**), suggesting a loss of selective network engagement during recall. Aged mice also displayed reduced hippocampal-amygdala ensemble coupling one week after training (**Figure 3H** and **Supp. Figure S7**), highlighting age-related connectivity deficits that may underlie memory impairments.

### Different neuronal types are part of hippocampal engrams

To better understand the neuronal composition of engram cells and their specific transcriptional response during memory retrieval, we performed snRNA-seq on sorted SUN1-GFP-tagged hippocampal nuclei from mice subjected to CFC, with or without recall (R vs. NR) seven days post-training (**Figure 4A** and **Supp. Figure S8A**; **Supp. Table 5**). In total, we analyzed the transcriptomes of 45,278 TRAPed hippocampal nuclei and detected over 32,000 genes (**Supp. Figure S8B-C**).

**Figure 4.**
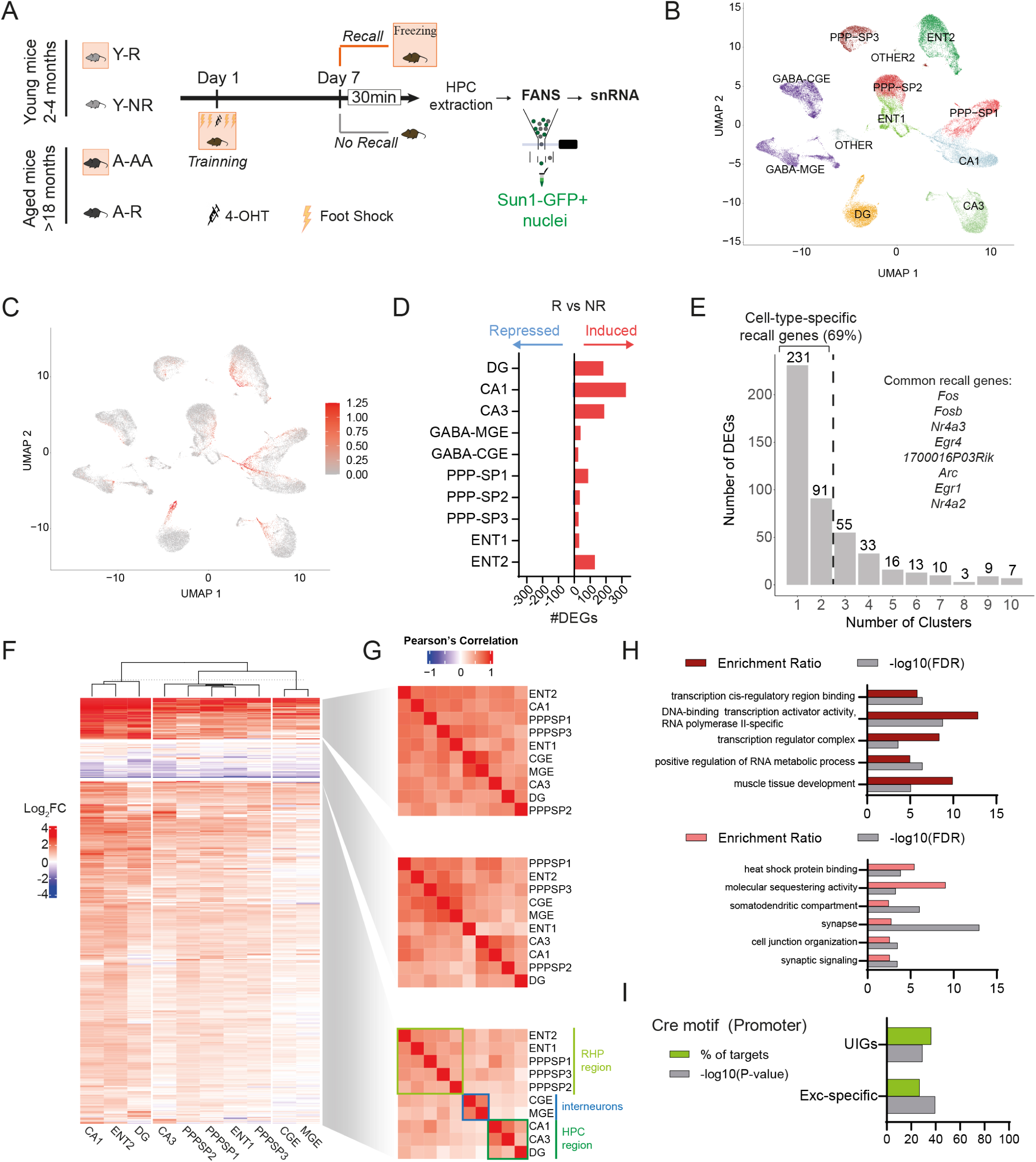
Memory retrieval evokes different transcriptional responses within hippocampal cells. **A.** Schematic representation of the experimental design. TRAP2 mice were initially exposed to CFC with a single dose of 4-oht labelling with SUN1-GFP cells activated in that moment. We prepare two behavioural cohorts: (i) no retrieval (NR) and (ii) memory retrieval (R), where mice were re-exposed to fear context. SUN1-GFP cells were sorted by flow cytometer and subjected to library preparation for single nuclei RNA sequencing (snRNA-seq). **B.** UMAP for snRNA-seq across hippocampus reveals distinct neuronal subtypes in the engram. **C.** UMAP plot showing module scores for 8 well known immediately early genes (IEGs) (*Fos*, *Nr4a1*, *Nr4a2*, *Nr4a3*, *Fosb*, *Npas4*, *Arc*, *1700016P03Rik*). **D.** DEGs for each neuron subtype in R versus NR of memory. **E.** Histogram of the number of neuron subtypes for which a recall DEG is significant. **F.** Heatmap of log 2-fold changes in DEGs for all neuronal types. DEGs were clustered by k-means. **G.** Pairwise correlation of the neuronal types for each cluster. **H.** GO analyses of UIG (red) and Memory-induced excitatory-specific genes (pink) clusters. **I.** Motif analysis for these two clusters.

To annotate the clusters, we applied the single-cell atlas created by Yao et al.,(*19*) (**Supp. Figure S8D**). This analysis identified 10 distinct hippocampal cell types activated during CFC (**Figure 4B** and **Supp. Figure S8E-G**). Memory retrieval did not change the cell-type composition within these clusters (**Supp. Figure S8F**). Consistent with prior studies in the amygdala or prefrontal cortex(*20*, *21*), the great majority of TRAPed cells were neurons, as evidenced by the expression of the neuronal marker NeuN (**Supp. Figure S8H-I**). Microglia and astrocytes were not detected among the TRAPed cells (**Supp. Figure S8J**). Furthermore, the expression of *Slc17a7*, a gene encoding the VGlut1 protein critical for glutamate transport, indicated that most of these neurons were excitatory (**Supp. Figure S8H-I**). These neurons originated from various hippocampal regions, including the CA1 (*Gm10754*), CA3 (*Il16*), and DG (*Prox1*) subfields (**Supp. Figure S9A-B**), and from retrohippocampal areas, such as the subicular complex and entorhinal cortex. Neurons from the subicular complex, including the parasubiculum, postsubiculum, presubiculum, subiculum, and prosubiculum, formed three distinct clusters (PPP-SP1-3) with region-specific marker expression (**Supp. Figure S9C-D**). Similarly, neurons from the entorhinal cortex were grouped into two clusters (ENT1-2), which exhibited markers for different cortical layers (**Supp. Figure S9E-F**). The integration of our snRNA-seq dataset with spatial transcriptomic data from the hippocampus(*22*) confirmed the anatomical assignment of the different clusters (**Supp. Figure S9G**).

Some inhibitory neurons were also tagged following CFC, which were classified into two major groups based on their developmental origins: medial ganglionic eminence (MGE) and caudal ganglionic eminence (CGE) (**Figure 4B**). Each group was further subdivided into three subclasses: CGE neurons were identified by markers such as *Lamp5*, *Sncg*, and *Vip*, while MGE neurons were marked by *Sst Chodl*, *Sst*, and *Pvalb* (**Supp. Figure S9H-J**). Interestingly, CGE-derived interneurons expressed *Prox1* (**Supp. Figure S9B**), a gene that promotes the maturation of this interneuron subtype(*23*).

### Memory recall evokes cell type-specific transcriptional signatures

To identify genes specifically involved in memory retrieval, we compared the R and NR conditions across different clusters of TRAPed cells. UMAP analysis revealed the displacement of several subsets of excitatory nuclei in the retrieval condition (**Supp. Figure S10A**). These nuclei were enriched for immediate early genes (IEGs) (**Figure 4C**), suggesting that the dominant source of variation between excitatory cells was related to their activation state. Although CA1, CA3, and PPP-SP1 neurons exhibited higher baseline expression of IEGs than other cell types, retrieval significantly increased the number of activated neurons across all cell types (**Supp. Figure S10B-C**). Using *Augur*(*24*), we observed similar responsiveness to retrieval across all neuronal types (**Supp. Figure S10D**). However, after pseudobulk analysis, principal neurons from the DG, CA1, and CA3 regions displayed the highest number of differentially expressed genes (DEGs) (*p* < 0.05) (**Figure 4D and Supp. Table 6**). To further explore neuronal interactions, we applied *NeuronChat* to infer cell-cell communication using our snRNA-seq data(*25*). Notably, intercellular communication involving granule neurons was enhanced in the retrieval condition (**Supp. Figure S10E**), highlighting their critical role in memory recall.

Interestingly, most DEGs (∼70%) were found in only one or two neuronal types (**Figure 4E**), suggesting that distinct gene sets were induced within each neuronal subtype. PCA revealed clear segregation of neuronal types based on their region of origin, clustering them into three groups: hippocampal, retrohippocampal and inhibitory neurons (**Supp. Figure S10F**). Many DEGs were specific to these broader categories (**Supp. Figure S10G**), indicating that activity-driven transcriptional programs are shaped by cellular ontogeny. DEGs analysis for each neuronal subtype indicated a general induction of memory-relevant genes (**Figure 4F**). Decision tree clustering grouped DEGs into three distinct categories (**Supp. Table 7**): i) ubiquitously induced genes (UIG) (n = 46), which were strongly upregulated across all neuronal subtypes, including well-known IEGs such as *Fos*, *Nr4a2* and *Arc*; ii) repressed genes (n = 34), which were slightly downregulated in some or all neuronal subtypes following memory recall; and iii) excitatory neuron-specific memory-induced genes (n = 386), which were upregulated in excitatory neurons but not in inhibitory neurons. This last group included genes known to be induced during the second wave of neuronal activation, such as *Nptx2*. A strong correlation was observed across all neuronal subtypes between the expression of UIGs and repressed genes (**Figure 4G**). Gene ontology (GO) analysis revealed that the UIG cluster was enriched for terms related to transcriptional activity, while the excitatory neuron-specific memory-induced gene cluster was enriched for terms related to synaptic plasticity (**Figure 4H**). Interestingly, both clusters of upregulated genes were enriched for CRE motifs (**Figure 4I**), suggesting CREB-mediated regulation of these transcriptional programs. No significantly enriched GO terms were identified for the repressed gene cluster.

Together, these findings suggest that memory retrieval engages distinct, cell type-specific transcriptional programs, with excitatory neurons exhibiting unique molecular signatures that may underlie their specialized roles in memory processing.

### Recall triggers distinct transcriptional responses in young and aged mice

To investigate the molecular basis of the dysfunctional activation observed in aged mice during memory retrieval, we conducted the same snRNA-seq screen (R vs. NR comparison) on 20-month-old mice. Merging data from young and aged mice, we analyzed the transcriptome of 74,268 hippocampal TRAPed cells, encompassing a total of 32,285 genes. Transcriptomic data from cells in aged mice displayed comparable quality and depth to those in young adult mice (**Supp. Figure S11A-B**). TRAPed cells were distributed in the same clusters in young and aged mice (**Figure 5A, Supp Fig S11C**) with no significant differences in the number of cells within each cluster (**Supp. Figure S11D**-**E**). As in young adult mice, no differences in cluster composition were observed between R and NR samples in the aged cohort (**Supp. Figure S11F**).

**Figure 5.**
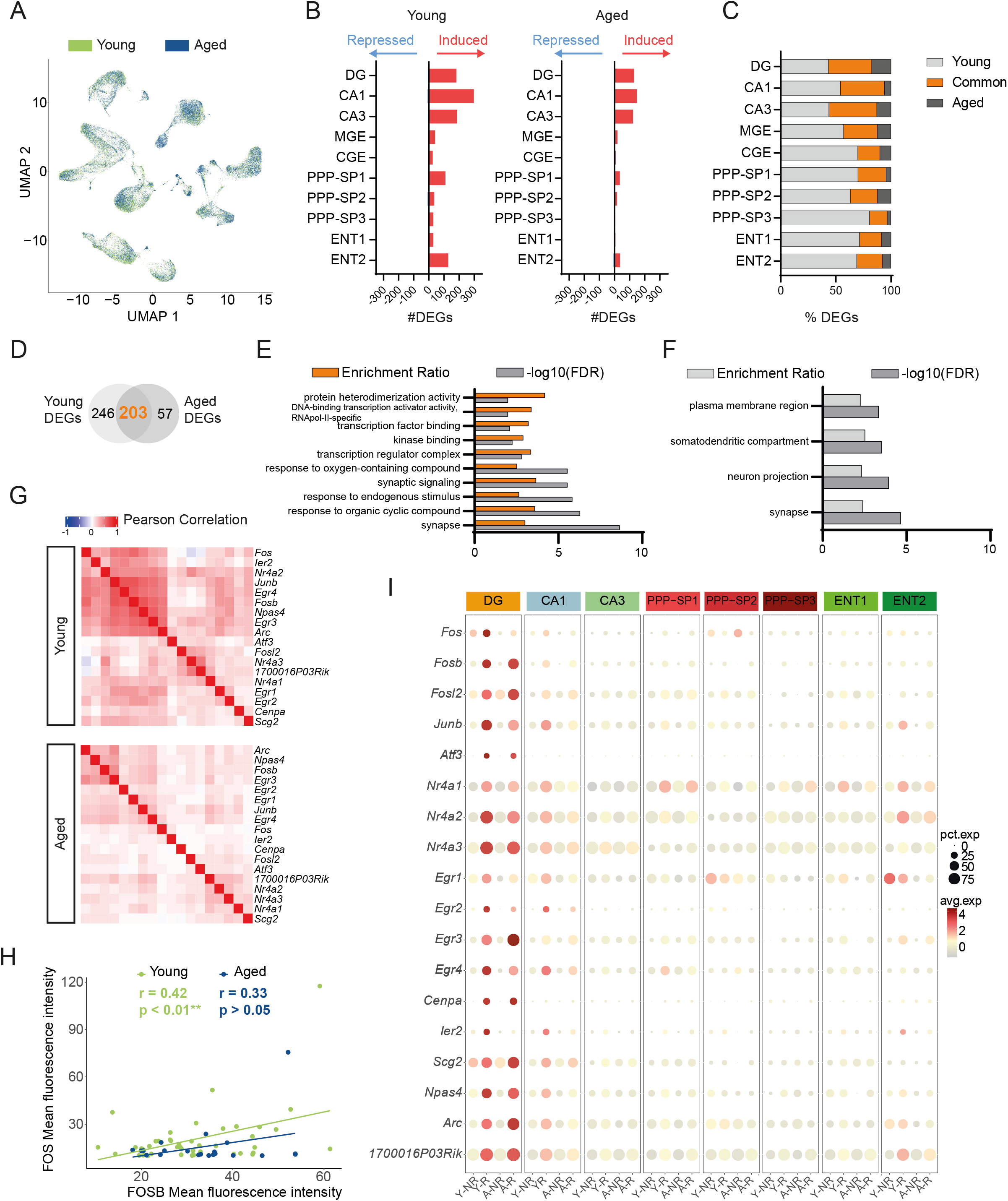
Recall produces distinctive transcriptional responses in young and aged mice. **A.** UMAP plot of analysis of snRNA-seq datasets from the SUN1-GFP cells of young (green) and aged (blue) mice. **B.** Number of DEGs for each neuron subtype in the comparison between R and NR for young (left) and aged (right) mice. Genes up- and downregulated in R condition shown in red and blue bars, respectively. **C.** Percentage of DEGs shared or age-specific for each neuronal type. **D.** Number of total DEGs shared or age-specific. **E.** GO significant terms for common DEGs between young and aged mice in R vs NR comparison. **F.** GO significant terms for young-specific DEGs. **G.** Pearson pairwise correlation of IEGs induction in strongly activated DG excitatory neurons (IEGs expression sum > 2) upon memory retrieval in young (top) and aged (bottom) mice. **H.** Pearson correlation between mean fluorescence intensity (MFI) obtained by immunohistochemistry against FOS and FOSB in young (orange) and aged (brown) mice for reactivated cells in DG. Each point represents a single nuclei SUN1-GFP+ FOS+ (MFI for FOS > 10). **I.** Bubble plot showing induction of major IEGs accross different types of activated excitatory neurons. Color reflects average normalized expression, while circle size indicates the percentage of IEG-positive nuclei across conditions (Y-NR: young no recall; Y-R: young recall; A-NR: aged no recall; A-R: aged recall).

Next, we examined whether fear recall triggered distinct transcriptional responses in neurons from aged mice (**Supp. Table 8**). A weaker induction of DEGs was observed in aged mice across all cell types (**Figure 5B**). In each cluster, approximately 50% of DEGs were unique to young mice (**Figure 5C**). The common DEGs (**Figure 5D**) were enriched for genes linked to transcriptional activation and synaptic plasticity (**Figure 5E**), while young-specific DEGs were primarily related to synaptic plasticity alone (**Figure 5F**), suggesting that second-wave genes are most sensitive to aging. The small set of aged-specific DEGs showed no enrichment for any biological function based on GO analysis. Motif analysis revealed significant enrichment of the CRE motif in both common and young-specific DEGs (**Supp. Figure S11G**)(*26*).

Interestingly, DG neurons exhibited the highest responsiveness to recall in aged mice (**Supp. Figure S12A**), differing from the distributed pattern observed in younger mice. A more detailed examination revealed enhanced induction of certain IEGs specifically in granule neurons of aged mice (**Supp. Figure S12B**), while younger animals displayed a more tightly coupled transcriptional response (**Figure 5G-H**). This pattern became more evident when restricting the analysis to highly activated nuclei (**Figure 5I**), indicating that granule neurons of aged mice display more widespread and less coordinated expression of IEGs (**Supp. Figure S12C-D**). These results align with the generalized fear response observed in aged mice.

We also assessed IEG expression in inhibitory neurons but did not identify subtype-specific changes, except for a slightly higher induction of IEGs in VIP interneurons of young mice (**Supp. Figure S12E-F**). This suggests that while excitatory neurons in aged mice undergo distinct transcriptional adaptations during memory recall, inhibitory neuronal responses remain largely unchanged.

### Memory recall induces a specific transcriptional program in reactivated cells

Although pseudobulk differential expression analysis in snRNA-seq experiments successfully identified changes in gene expression associated with recall by comparing the subset of reactivated neurons within each cluster of TRAPed cells, these analyses could not address the differences between reactivated neurons and other neurons activated during retrieval. To explore in greater detail and coverage the transcriptional signature of reactivated neurons, we focused on DG neurons because they displayed the largest age-dependent difference during reactivation and their critical role in context discrimination. NucTRAP2 mice were subjected to CFC (**Figure 6A**) and sacrificed one week after training and 90 minutes after recall, DGs were micro-dissected and dissociated, and three different populations of granule neurons were isolated using FANS, based on GFP fluorescence and FOS immunostaining (**Supp. Figure S13A-E**): (i) green cells (SUN1-GFP^+^ neurons), activated only during fear training (1^st^ experience cells), (ii) red cells (FOS^+^ neurons) activated only during recall (2^nd^ experience cells), and (iii) yellow cells (SUN1-GFP^+^/FOS^+^ neurons) activated both during memory encoding and recall (reactivated cells). A total of 100 nuclei from each population were collected from the DG of four mice and used to generate libraries for nuclear RNA-seq (nuRNA-seq) with ultra-low input. To validate the sorted populations, in silico analysis confirmed their identity: SUN1-GFP expression was observed in 1^st^ experience and *reactivated* cells (**Supp. Figure S14A**), while increased IEG expression was detected in 2^nd^ experience and reactivated cells (**Supp. Figure S14B**). PCA showed the clustering of replicates for each population, with the transcriptional profile of *reactivated* cells positioned between those of the other two populations (**Figure 6B**).

**Figure 6.**
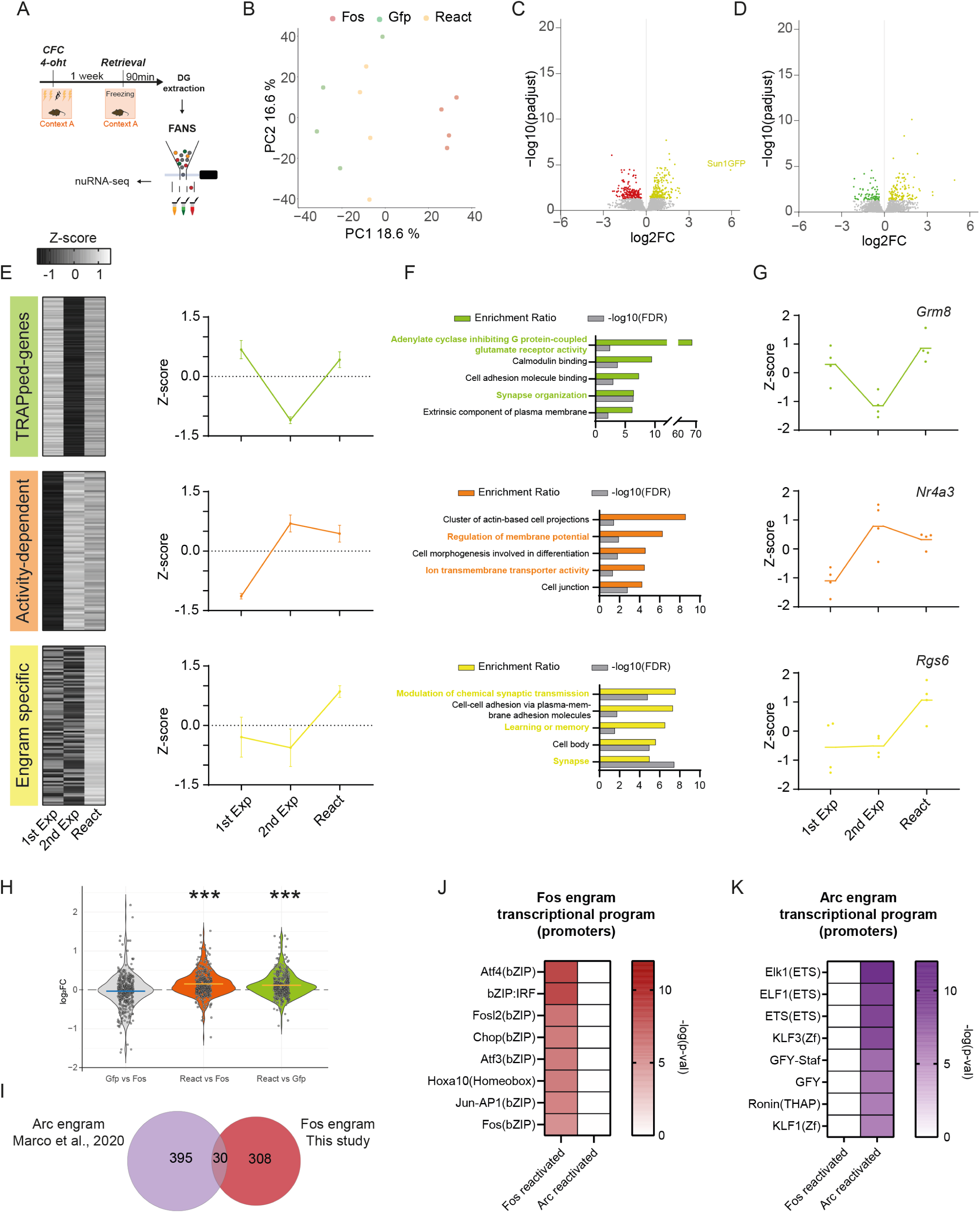
Transcriptional programmes activated by memory retrieval in reactivated cells are distinct to new activated cells. **A.** Schematic representation of experimental design. **B.** PCA for each neuronal population sorted. **C.** Volcano plot showing the significance and p-adjust distribution after differential gene expression analysis between Reactivated and 2^nd^ experience cells. **D.** Volcano plot showing the significance and p-adjust distribution after differential gene expression analysis between Reactivated and 1^st^ experience cells. **E.** Heat map of z-scores of normalized counts and mean z-score of each population by cluster. **F.** GO terms for each cluster. **G.** Representative DEGs for each cluster. **H.** Comparison between three sorted populations of the expression levels of the Arc-engram specific genes obtained from Marco et al. (2020)(*28*) (one sample t-test: \*\*\**P*<0.001). **I.** Venn diagrams showing the overlap between upregulated genes in reactivated cells of Fos- and Arc- engrams. **J-K.** Motif analysis of upregulated genes in Fos- (J) and Arc- engram (K).

A total of 339 DEGs were found to be upregulated in reactivated cells (**Figure 6C-D**) and clustered based on their expression patterns (**Figure 6E**): (i) 162 genes highly expressed in both 1^st^ experience and reactivated cells, termed “TRAPed genes”; (ii) 85 genes upregulated in recently activated populations (2^nd^ experience and reactivated cells), termed “activity-dependent genes”; and (iii) 92 genes uniquely enriched in reactivated cells, referred to as “engram-specific” genes. GO analysis was conducted for each cluster (**Figure 6F**). The TRAPed cluster was enriched for genes involved in “synapse organization” and “inhibitory mechanisms of glutamate receptors”, such as *Grm8* (**Figure 6G**), suggesting that persistent upregulation of these genes support memory consolidation(*20*). The activity-dependent cluster was enriched in terms related to signal transduction and cellular activation, including IEGs such as *Nr4a3* (**Figure 6G**). Finally, the engram-specific cluster showed enrichment for terms associated with learning, memory and synaptic plasticity. This cluster included genes such as *Rgs6* (**Figure 6G**), which has been previously implicated in memory processes in the DG(*27*). Notably, while the activity-dependent cluster was associated with postsynaptic terms, the engram-specific cluster was more closely linked to presynaptic functions (**Supp. Figure 14D**). This distinction may be related to the increased outputs from DG during memory retrieval observed in the snRNA-seq analysis (**Supp. Figure 10F**).

To investigate the specific contributions of inhibitory and excitatory synaptic plasticity during recall, we compared the normalized counts of canonical markers from each population. As expected, non-neural markers were minimally expressed, while neuronal markers, such as *Rbfox3*, were highly expressed (**Supp. Figure 14E**). The expression of inhibitory neuronal markers was detected in the three neuronal populations, although intriguingly somatostatin (*Sst*) interneurons (belonging to the GABA-MGE group) were slightly enriched in reactivated cells (**Supp. Figure 14F**), accompanied by a relative depletion of excitatory markers such as *Prox1* (**Supp. Figure 14E**). These results were confirmed through the detection of reactivated somatostatin interneurons in the hilus by immunohistochemistry (**Supp. Figure 14G**).

Finally, we compared our results with those of the only previous study investigating the transcriptional signature of reactivated cells, conducted in ArcTRAP(1) mice(*28*). We observed that 438 DEGs associated with reactivation during memory recall in the Arc-dependent engram(*28*) were also upregulated in the Fos-dependent engram (**Figure 6H**). However, the overlap between DEGs upregulated in reactivated cells from both studies was relatively low, at approximately 10% (**Figure 6I**). Interestingly, while DEGs in the Fos-dependent engram were enriched for genes containing AP1 binding sites (**Figure 6J**), DEGs in the Arc-dependent engram were enriched for genes bound by ETS TFs (**Figure 6K**). These results are consistent with the limited overlap between Arc and Fos-expressing granule cells after salient experience(*29*). Together, these findings suggest the coexistence of distinct engrams in the hippocampus, potentially encoding different aspects of memory.

### Aging attenuates the transcriptional signature of reactivated cells and exacerbates GABAergic connectivity

To assess whether genes upregulated in reactivated cells during recall are affected in aged mice, we integrated nuRNA-seq data from DG micro-dissections with snRNA-seq data corresponding to three neuronal types in the DG region (DG granule neurons, GABA-MGE, and GABA-CGE clusters). This analysis revealed that young mice exhibited higher expression of genes upregulated in reactivated cells during retrieval in the DG and GABA-MGE clusters, but not in GABA-CGE (**Figure 7A–C**). Significant positive correlations between gene expression from nuRNA-seq (fold change in reactivated vs. 1^st^ experience cells comparison) and snRNA-seq (fold changes in activated vs. non-activated cells comparison) were observed in the DG and GABA-MGE clusters of young mice (**Supp. Figure 15A**). However, in aged mice, significant positive correlations were found only in the DG cluster. These findings suggest impaired engram reactivation in DG and GABA-MGE neurons in aged mice.

**Figure 7:**
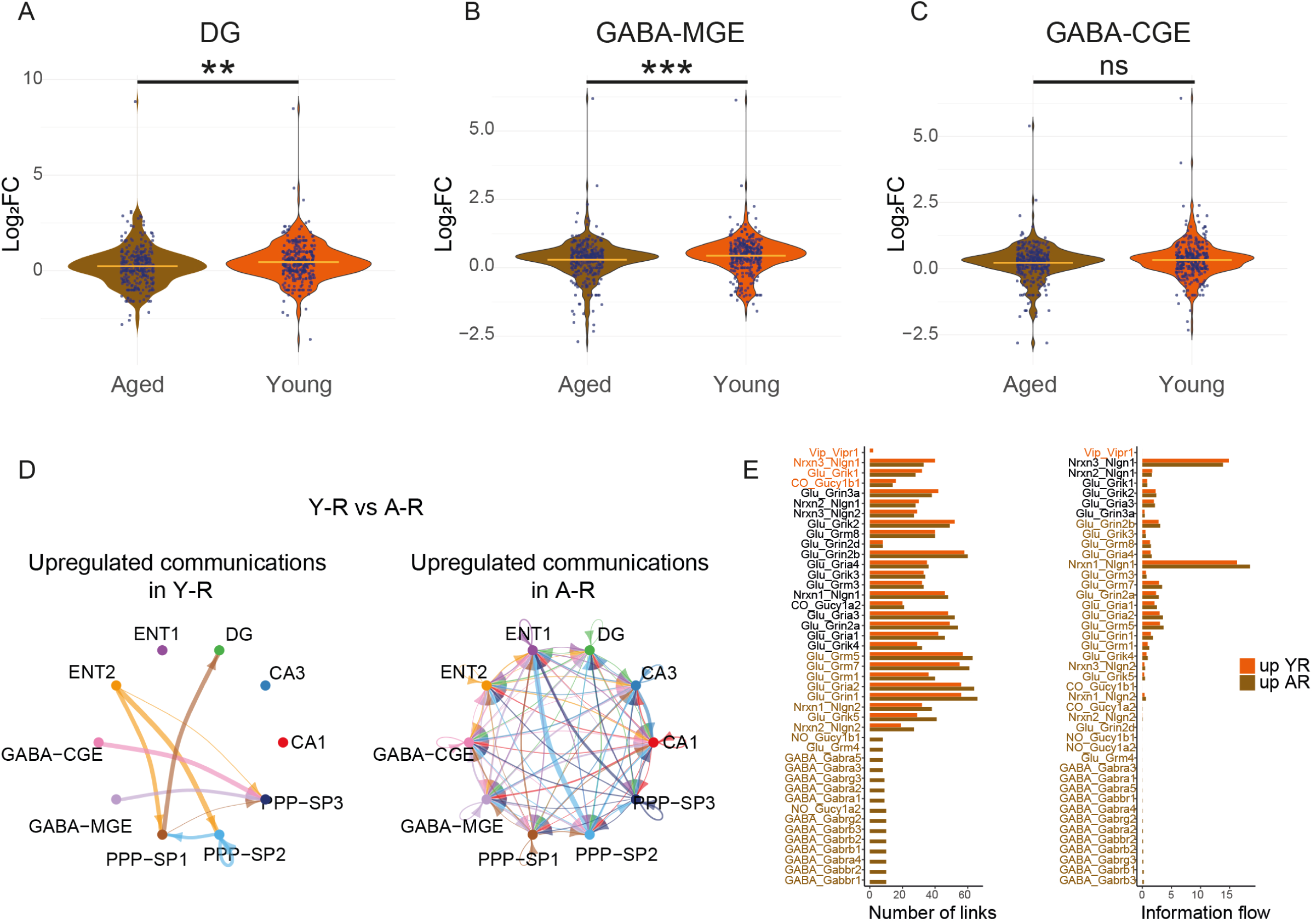
Weaker transcriptional response of engram related genes in aged mice during memory retrieval. A-C. Comparison between young and aged mice of expression level of upregulated genes in reactivated cells from nuRNA-seq for DG (A), GABA_MGE (B) and GABA-CGE (C) clusters. **D.** Circle plots from NeuronChat showing upregulated intercellular communication in Y-R (left) and A-R (right). The width of each link indicates the absolute difference between Y-R and A-R, in the sum of communication strength values over all interaction pairs. **E.** Bar plot comparing the number of links (left) and informational flow (right) between Y-R and A-R for each ligand-receptor pair.

To investigate whether connectivity in these clusters was altered with age, we applied *NeuronChat* to the retrieval group for each age. Aged mice exhibited increased intercellular communication (**Figure 7D, Supp. Figure 15B**), particularly, through upregulated GABAergic connections (**Figure 7E**). Notably, interactions such as GABA-Gabbr1, GABA-Gabra4, and GABA-Gabrb1 showed robust upregulation in aged mice, with outputs from GABA-MGE cells targeting all other cell types (**Supp. Figure 15C-D**). Among these GABA-MGE cells, somatostatin-expressing interneurons have been suggested to negatively regulate the size of neuronal ensembles(*30*), which could explain the reduced DG engram size during context-specific recall observed in aged mice (**Figure 3D**). In contrast, consistent with previous results (**Supp. Figure 12C**), only VIP interneuron-related connectivity was upregulated following CFC in young mice (**Figure 7E**). These results suggest that increased inhibition, driven by dysfunctional GABA-MGE neurons during memory retrieval, may impair engram reactivation in aged mice.

## Discussion

### Brain-wide mapping of CFC memory encoding and recall

Clearing techniques have been employed to map cells activated during fear conditioning using antibodies against FOS(*31*) or FosTRAP(1)-tdTomato tagging(*8*, *32*). Here, we combined the FosTRAP2 strain with the SUN1-GFP reporter line to provide a more sensitive and efficient labelling(*6–8*) and generate the most comprehensive mapping of engram cells across the mouse brain at different ages to date.

A previous study investigated the reactivation of a fear memory engram across the brain using FosTRAP(1) mice crossed with a tdTomato (tdT+) reporter(*32*). While this pioneering work was a major step toward brain-wide engram mapping, the high activation threshold for tagging and interference with endogenous *Fos* expression limited its ability to provide a complete picture of memory-related activation(*8*). Additionally, the use of tdT, which labels the entire cell, reduced the efficiency of automatic detection algorithms in counting individual cells. In comparison, our study employed a more refined approach by combining the higher sensitivity of the TRAP2 strain with the precise nuclear envelope labeling of SUN1-GFP. This strategy significantly improved the accuracy of individual cell quantification and enabled the creation of a more complete and precise map of a specific memory. Furthermore, we compared the formation of the CFC engram to control conditions, including scenarios where the memory did not form (HC) or where recall did not occur (A-B condition). Overall, we quantified cellular ensembles bearing a memory trace in 378 brain regions, providing a valuable reference to explore the distribution, functional connectivity, and age-related changes of memory engrams. This established our study as the most extensive engram mapping effort to date, shedding light on engram formation and stability across brain regions.

DeNardo et al. revealed distinct reactivation patterns for recent (1 day post-training) and remote (14 days) fear memory in the prelimbic cortex using TRAP2 mice(*8*). Expanding on this, we compared engram reactivation at one and three weeks in the entire brain. Notably, for the first time, we report a shift in engram connectivity across brain regions during memory consolidation: Fos ensembles showed stronger coupling between the hippocampus and amygdala at one week, while at three weeks, the correlation between amygdala and prefrontal cortex ensembles became more pronounced. These findings provide critical insights into the dynamic nature of memory engrams over time and lay the groundwork for future studies exploring how memory consolidation mechanisms evolve across different brain regions.

### Transcriptional signatures of CFC memory encoding and recall

We used snRNA-seq to analyze the transcriptional response to memory retrieval in a cell-type-specific manner. While canonical IEGs were shared across most cell types, most other genes induced during retrieval were cell-type-specific. Notably, neurons originating from anatomically closer regions exhibited more similar transcriptional responses. Cell type-specific genes induced during memory recall were associated with synaptic plasticity, potentially supporting the synaptic reorganization that occurs during memory reconsolidation(*33–35*). Furthermore, these genes were enriched for CRE motifs, underscoring the critical role of CREB in memory processes(*26*, *36*).

Additionally, the combination of snRNA-seq with low-input nuRNA-seq for 1^st^ experience (memory encoding), 2^nd^ experience (recall), and reactivated cells provided unprecedented insight into the transcriptional signatures associated with FOS-dependent engrams. Until now, it was unclear whether reactivated engram cells exhibit the same transcriptional response as cells activated for the first time to encode other aspects of an experience. Here, we identified a transcriptional response unique to reactivated DG granule neurons, characterized by the expression of a set of genes encoding key proteins involved in excitatory synapse assembly, synaptic plasticity and memory, such as *Rgs6*, *Ntrk2* and *Prkcg*. This underscores the distinct role of these cells in maintaining a memory trace.

Sun et al. provided causal evidence for functional heterogeneity in memory trace encoding(*37*), identifying two distinct engrams dependent on NPAS4 and FOS, which encode different aspects of memory. However, the transcriptional response specific to each type of engram had not been characterized. Leveraging findings from studies on *Arc*-dependent engrams during memory reactivation(*28*), we found that *Fos*- and *Arc*-dependent engrams are transcriptionally distinct. *Fos*-dependent engrams are marked by a transcriptional cascade related to AP-1, whereas Arc-dependent engrams are more closely associated with the ETS family of transcription factors, such as *Elk1*, which has also been implicated in synaptic plasticity and learning(*38*).

Overall, our findings provide critical insights into the molecular mechanisms underlying distinct memory engrams, significantly advancing our understanding of transcriptional heterogeneity in memory processing. Moreover, by detailing cell-type-specific transcriptional responses following retrieval, our study expands the current understanding of memory transcriptional programs and their role in engram stability.

### Age-dependent differences in CFC memory encoding and recall

While the spatial organization of neurons has been observed to remain stable with age(*39*), we identified age-dependent changes in functional engram connectivity between brain regions, as well as differences in the transcriptional signature for reactivation in specific neuronal types. These differences may contribute to age-related memory deficits.

Single-cell studies have provided new insights into the cellular and molecular biology of aging neurons, primarily focusing on the basal transcriptional profile and the role of non-neuronal cells and inflammatory responses(*40–42*). However, these studies have largely overlooked the impact of aging on neuronal transcriptional responses during learning. Our results demonstrate that while engram reactivation in aged and young mice shares similarities, it also exhibits notable differences. Although many genes related to transcriptional regulation were induced upon retrieval in both groups, significant differences emerged in the level of induction at the single-neuron level for certain IEGs, particularly transcription factors. These findings suggest dysfunctional engram activation in aging, especially in the DG. Furthermore, several genes associated with synaptic plasticity that were strongly induced in young mice were not significantly induced in aged mice. These findings suggest that age-related differences in cortical reorganization during memory consolidation(*15*) along with altered transcriptional responses of engram neurons in the hippocampus—a region crucial for integrating contextual cues during recall(*43*, *44*)—may contribute to memory deficits and less accurate retrieval in aging. Taken together, these findings advance our understanding of the molecular and cellular mechanisms underlying cognitive aging and help identify potential therapeutic targets for age-related cognitive decline.

## Methods

### Mouse strains and treatments

The generation of CamKIIα-CreERT2(*45*) (from the EMMA EM:02125), TRAP2 (pFos-CreERT2)(*7*, *8*) (from Jackson Lab stock #030323), CAG-SUN1/sfGFP mice(*46*) (from Jackson Lab stock #021039, RRID:IMSR_JAX:021039) and CAG-[STOP]-tdTomato(*13*) (from Jackson Lab stock # 007914) have been previously described. All mice were maintained on a C57BL/6 J genetic background. Mice were maintained and bred under standard conditions, consistent with Spanish and European regulations. All animal protocols were approved by the Animal Welfare Committee at the Instituto de Neurociencias, the CSIC Ethical Committee and the Dirección General de Agricultura, Ganadería y Pesca of Generalitat Valenciana. Mice were kept in a controlled environment with a constant temperature (23 °C) and humidity (40-60 %), on 12 h light/dark cycles, with food and water ad libitum. They were maintained in a sterile room located within the Animal House at the Instituto de Neurociencias (CSIC-UMH). CamKIIα-CreERT2 x CAG-SUN1/sfGFP mice x CAG-[STOP]-tdTomato mice were used to label excitatory neurons of the forebrain after 5 doses of TMX (Sigma Aldrich, 20 mg/mL dissolved in corn oil) administrated intragastrically(*47*). TRAP2 x CAG-SUN1/sfGFP and TRAP2 x CAG-SUN1/sfGFP x CAG-[STOP]-tdTomato were used to label permanently activated cells administrating 4-OHT (50 mg/kg) via intraperitoneal injection, this was prepared as previously described(*8*). The experiments were carried out on animals of different ages. Young mice were between 2 and 4 months old and aged mice were between 18 and 22 months old. For the induction of brain activity by KA (Milestone PharmTech USA Inc.), a concentration of 25 mg/kg was injected intraperitoneally.

### Novel exposure

For novel exposure (NE) mice received 2 days of 5 min handling before exposing them to a novel environment. It was conducted in 48 x 48 x 30 cm white acrylic glass boxes, where mice were allowed to freely move for 120 min. To evaluate the levels of labelling comparing heterozygous versus homozygous TRAP2-SUN1GFP mice or leakiness in these mice, 4-OHT or oil was administrated after 60 first minutes. Mice behaviour was monitored by the video tracking system SMART (Panlab S.L. Barcelona, Spain). To evaluate motility, total distance was estimated.

### Discriminative contextual fear conditioning

For the extended contextual fear conditioning (CFC), mice were subjected to a 120-minute-long training session in a 40 x 40 cm arena with objects and an electrified grid, termed Context A. They received two footshocks (0.5 mA, 2 sec) during the first 10 minutes, one each 5 min. 4-OHT was administrated 60 min after to start the training. When mice were replaced in context after 4-OHT administration, they received another two footshocks during the first 10 min of the second hour of context exposure (0.5 mA, 2 sec). After 7 days, mice were re-exposed to the same context during 15 min to evaluate long-term memory and perfused 90 min after of start the recall to evaluate engram formation.

To evaluate if the freezing is dependent of the context, other batch of mice were trained in Context A but they were placed in a different context, termed Context B, 7 days after training. Also, they were perfused 90 min after starting the test. Freezing score was calculated as the percentage of time for which the mice remained immobile. Immobility for more than 2 s was counted as freezing behaviour(*48*). For a more stringent test for context discrimination to confirm age-related memory imprecision. Mice were trained in classical CFC chamber (Panlab S.L., Barcelona, Spain), which was equipped with an electrified grid. In the training session, mice were allowed to explore the fear conditioning chamber for 2 min. Afterwards, they received a 0.5 mA, 2 sec footshock. Then, mice had 56 seconds for recovery and received an identical second footshock and mice remained in the box for 1 additional minute. The time animals remained still (freezing) was registered through a piezoelectric sensor located at the bottom of the fear box. To assess contextual memory, seven days later after the conditioning session mice were returned to the same box for 3 min, referred as Context A’, and their freezing behaviour was measured, three hours after recall context were exposed to a different one, referred as Context B’, to obtain discrimination index (DI). Finally, eight days after training mice were re-exposed to same context exposure than day seven but in this case beginning by Context B’.

### Immunohistochemistry and microscopy

Mice were anesthetized with an injection of ketamine/xylazine, perfused with 4% paraformaldehyde (PFA) in PBS and postfixed overnight at 4 °C. Coronal and sagittal vibratome sections (50 μm) were obtained. These were washed in PBS and PBS − 0.1% Triton X-100 (PBT) and incubated for 1 h at room temperature with blocking buffer 4% newborn calf serum (NCS, Sigma N4762) in PBT. After, sections were incubated overnight at 4 °C with the primary antibodies diluted in 5% NCS-PBT. The primary antibodies used were: α-Fos (1:500; Thermo scientific, T.142.5), α-GFP (1:1,000; Aves Labs, GFP-1020), α-DsRed (1:1,000; Clontech, 632496), α-Iba1 (1:500; Wako, 019-19741), α-GFAP (1:500; Sigma, G9269), α-Cre (1:500; Sigma, C7988), α-Sst (1:100; ImmunoStar, 20067). After primary antibody incubation and washing with PBS, sections were incubated with fluorophore-coupled secondary antibodies which were acquired from Invitrogen and used in a dilution 1:500. Sections were counterstained with a 1 nM DAPI solution (Invitrogen). Images were taken using a Leica SPEII vertical confocal microscope and processed using Fiji-ImageJ software with the *Cell Counter* plugin. For the analysis of FOS and FOSB fluorescence intensity, the mean gray value was measured for each SUN1⁺ nucleus. Nuclei were considered FOS⁺ when their mean intensity exceeded a threshold value of 10.

### iDISCO+ Sample Processing

Right after the behavioural paradigm, mice were deeply anesthetized (ketamine/xylazine) and perfused with 4% paraformaldehyde (PFA). The brains were collected and postfixed by immersion in 4% PFA for 3h at room temperature (RT). Then they were washed in PBS 1X at RT. For preservation, brains were kept in PBS 1X with 0.01% Sodium Azide at 4°C.

Whole brain immunostaining and clearing was performed following the iDISCO^+^ protocol(*12*) with minimal modifications. All the steps of the protocol were done at room temperature with gentle shaking unless otherwise specified. All the buffers were supplemented with 0,01% Sodium Azide (Sigma-Aldrich, Germany) to prevent bacterial and fungal growth. Fixed brains were dehydrated in a methanol dilution series. Aqueous solutions of methanol (Sigma-Aldrich, France) at 20%, 40%, 60% and 80% were used for 1 hour incubations, followed by two incubations in 100% methanol (one and two hours respectively). After dehydration, the samples were placed in a solution of 6% hydrogen peroxide (Sigma-Aldrich) in methanol, and bleached overnight at 4°C without shaking. In the second day, the samples were rehydrated by sequential incubations in 60%, 40%, and 20% methanol solutions for 1 hour each. After methanol pretreatment, the samples were washed twice in PBS for 15 minutes and then permeabilizated by an incubation in PBS containing 0.2% Triton X-100 (Sigma-Aldrich) for one hour, followed by further permeabilization through an overnight incubation at 37°C in Permeabilization Solution (composed by 20% dimethyl sulfoxide (Sigma-Aldrich), 2,3% Glycine (Sigma-Aldrich, USA) in PBS-T). After permeabilization, the samples were blocked in a solution of 0.2% gelatin (Sigma-Aldrich) in PBS-T for 24 hours at 37°C. The same blocking solution was used for antibody incubations. Primary antibodies were incubated for 10 days at 37°C with gentle shaking, using Fos (1:1000; Synaptic Systems, #226 017) and GFP (1:2000; Aves Labs, GFP1020) as primary antibodies. The samples were then washed in PBS containing a 0.2% of Tween 20 and 1ug/mL of Heparin (PBSTwH) twice for 1 hour and a third wash overnight. After washing the primary antibodies, the samples were incubated for 8 days with secondary antibodies (Donkey anti-Chicken, 1:1000, Thermo scientific #A78949; and Donkey anti-Rabbit, 1:1000, Thermo scientific, #A31573). After immunostaining, the samples were washed in PBSTwH (two washes of 1 hour and a third wash overnight) and then dehydrated through a gradient of methanol/water solutions (20%, 40%, 60%, 80%, and 100%, each for 1 hour), finishing with an overnight incubation in 100% methanol. For the final clarifications, the samples were incubated 3 hours in a solution of 66% dichloromethane and 33% methanol, then washed twice in 100% dichloromethane (15 minutes each), and immersed in dibenzyl ether (Sigma-Aldrich) until light sheet imaging.

### Light-Sheet Imaging

An Ultramicroscope Blaze (Miltenyi Inc) was used for whole hemisphere image acquisition. Two types of images were done: the specific signal of GFP and Fos immunolabeling, and an anatomical overview of the brain.

To image the immunolabeling signal, a 4X MI PLAN objective lens with 0.35NA was used. GFP immunolabeling (revealed with Alexa 555) was acquired using a laser line 561nm (100 mW) and a 595/40 filter. FOS immunolabeling (revealed with Alexa 647) was acquired using a laser line 639nm (70mW) and a 680/30 filter. The field of view was cropped to 1000×1000 pixels, and a mosaique acquisition was set, covering the whole hemisphere with a 5% of overlap in between tiles. To optimize the spatial equivalence of the 3D acquisitions done for each channel (GFP and FOS), the laser/filter change was done at every z-step. The laser power and exposure time were kept constants for all the acquisitions. To obtain an overview of the anatomy of the brain that enabled signal alignment to a reference atlas, the intrinsic autofluorescence of the sample was captured. A 1.1X MI PLAN objective lens (0.1NA) was used in combination with a 488nm laser (85mW) and a 525/50 filter. The field of view was cropped to fit the size of the whole hemisphere. For consistency, the right hemisphere of all the animals was imaged in sagittal orientation. All the acquisitions were done with triple laser illumination from the left side, with a laser numerical aperture of 0.03, maximal laser width and a step size of 5 µm. The microscope was equipped with a 4.2 Megapixel camera (sCMOS sensor), obtaining a lateral resolution of 1.62 µm/pixel with the 4X objective lens, and 5.91 µm/pixel with the 1.1X lens.

### ClearMap2 FOS+ and SUN1-GFP+ cell counting

The images obtained with the lightsheet microscope were analyzed with ClearMap 2(*11*, *12*). Mosaique acquisitions obtained for GFP and FOS were stitched and processed for background subtraction and local maxima detection. A watershed was generated around the local maxima considering a minimal intensity threshold. Finally, a volume filter was applied to select only those extended maxima with a size compatible with the cell nucleus (expected for GFP and FOS stained). The stitched image for FOS and GFP was then aligned to a reference atlas (Allen Brain Atlas, 2022), using elastix (https://elastix.lumc.nl). The autofluorescence scan was used as internal anatomical template for each sample. The coordinate transformations obtained were applied to every maxima detected, generating density maps in the common coordinate system of the Allen Brain Atlas, and a data frame file with the counts of detected cells per brain region.

### Fluorescence-activated nuclei sorting (FANS)

All steps were performed at 4 °C unless indicated otherwise. DGs or whole hippocampus was processed. Hippocampi were microdissected after cervical dislocation and stored at −80 °C. DGs were dissected after cervical dislocation as has been previously described in other study (Hagihara et al., 2009) and processed immediately. Nuclei were extracted by mechanical homogenization using a Dounce homogenizer (Sigma) with 500 µL of nuclear extraction buffer (NEB: Sucrose 250 mM, KCl 25 mM, MgCl2 5 mM, HEPES-KOH 20 mM (pH 7.8), IGEPAL CA-630 0.5 %, Spermine 0.2 mM, Spermidine 0.5 mM, and 1x proteinase inhibitors (cOmplete EDTA-free, Roche), 60 U ml–1 RNasin Plus RNasa (Promega, N2611)). For nuRNA-seq experiment, samples were filtered using a 35 μm nylon mesh-capped tube, and the samples were filled up to 750 µL with NEB. Nuclei were transferred to a 1.5 mL eppendorf tube and incubated for 10 minutes on a rotator with blocking buffer containing 3% newborn calf serum (NCS, Sigma N4762). After incubation, the Eppendorf tubes were centrifuged at 1500 G for 5 minutes. The supernatant was discarded, and the nuclei were resuspended in 500 µL of NEB. For nuclear staining, nuclei were incubated for 30 min with the primary antibody α-Fos (1:500; Thermo scientific, T.142.5) and 3% NCS on a rotator. Then, they were centrifugated at 1500 G for 5 min and supernatant was discarded. Nuclei were resuspended again in 500 µL of NEB. Secondary antibody (1:1,000) with 3% NCS and DAPI (at a final concentration of 0.01 mM) were incubated for another 15 min on a rotator, centrifugated again at 1500 G for 5 min and resuspended in 500 µL of PBS 1X. Nuclei populations were selected using a FACS Aria III (BD Biosciences) for DAPI staining, singlet nuclei, SUN1-GFP+ and FOS+ positive staining, and sorted into three different eppendorf (one for each population). Flow cytometry data were analysed and plotted using FlowJo (v.10). For snRNA-seq, samples were filtered using a 35 μm nylon mesh-capped tube and filled up to 750 µL with NEB. These steps were repeated for one or two mice of the same condition, and the nuclei from the mice were pooled into the same tube by condition. A total of 1.5 mL of PBS supplemented with additives (0.2 mM Spermine, 0.5 mM Spermidine, 1x proteinase inhibitors [cOmplete EDTA-free, Roche], 0.2 U µL⁻¹ RNasin Plus RNase inhibitor [Promega, N2611], and 1% bovine serum albumin) was added to the tube containing the pooled nuclei. SUN1-GFP+ nuclei were selected using a FACS Aria III (BD Biosciences) by gating for singlet nuclei and SUN1-GFP+ positive fluorescence and were sorted into a single Eppendorf tube. Flow cytometry data were analysed and visualized using FlowJo (v.10).

### Nuclear RNA-seq: Library preparation and sequencing

After FANS, 100 hundred nuclei for each population by mouse were obtained. RNA-Seq libraries were generated with a SMARTer Stranded Total RNA-Seq Kit v3 - Pico Input Mammalian (Takara Bio Inc., Shiga, Japan), according to the manufacturer’s instruction. Total RNA was converted to cDNA and the cDNA derived from rRNA was depleted by ZapR v3 & R-Probe v3 contained in SMARTer Kit during the procedure. Adapters with UMIs are included to remove PCR duplicates. For sequencing, paired-end 100 bp was conducted in a NextSeq 2000 system (Illumina) and a depth of at least 50 million reads per sample.

### Data analysis

To perform quality control of sequenced reads we used FastQC (version 0.11.9). To extract unique molecular identifiers (UMIs), removed PCR duplicates and retained only reads that represent individual RNA molecules for quantification we used UMI-Tools(*49*) (version 1.1.2). Reads were trimmed with TrimGalore (version 0.6.7; https://www.bioinformatics.babraham.ac.uk/projects/trim_galore/) to remove adapter sequences and STAR(*50*) (version 2.7.9) was used to align reads to the mouse genome (GRCm38, mm10), reads with a mapq > 30 were selected using Samtools (version 1.13). For the evaluation of SUN1-GFP expression, all nuRNA-seq samples were used to obtain SUN1-GFP sequence of ROSA26 using Trinity (version 2.15.1) and it was introduced as an additional chromosome into the mouse reference genome. Counts were quantified to gene using Rsubread (version 2.12.3) and mouse genome (GRCm38, mm10) with SUN1-GFP sequence introduced. Differential expression analysis was performed using Deseq2 (version 1.38.3). Wald test was used to obtain DEGs considering p-adjusted < 0.05. DEGs of reactivated cells in either of the two comparisons (Reactivated vs FOS and Reactivated vs SUN1-GFP) were classified into three distinct groups based on the SUN1-GFP vs FOS comparison. DEGs from reactivated cells that were significantly upregulated in SUN1-GFP were assigned to the TRAPed-genes group, while those significantly upregulated in FOS were categorized as Activity-dependent genes. For the visualization of DEGs derived from nuRNA-seq data, Z-scores for each gene were calculated based on the normalized counts from the respective samples.

Gene ontology analysis, *Biological process*, *Cellular component* and *Molecular function* were performed using WebGestalt (WEB-based GEne SeT AnaLysis Toolkit(*51*); www.webgestalt.org), using over-representation analysis (ORA) as the enrichment method. Motif analysis of DEGs was performed using HOMER (version 4.11.1). In addition, we have performed SynGO(*52*) (version 1.2) analysis with default parameters, using brain expressed genes as background.

### Single nuclei RNA-seq: Library preparation and sequencing

After FANS, 20,000 nuclei per sample (pool of 2-3 animals) were loaded into the single cell Chromium Next GEM Chip G and then the generation of barcode-containing partitions was carried out with the Chromium Controller (10X Genomics). Chromium Next GEM Single Cell 3’ Library (v3.1) was employed for post-GEM-RT clean-up, cDNA amplification and the generation of barcoded (Dual Index Plate TT Set A) libraries. Libraries were sequenced to an average depth of 50K reads per nuclei on a NovaSeq X Plus Series PE150 sequencer.

### Data analysis

Sequenced samples in FASTQ format were analyzed using Cell Ranger (10X Genomics, version 7.1.0) and aligned to the CRGm38 (mm10) mouse reference genome. Output data were pre-processed and analysed using Seurat R package (version 5.0.0). Low-quality cells were filtered out (genes detected < 1000 or > 7000, number of counts < 800 and mitochondrial genes > 5%) and doublets were detected and removed using R package scDblFinder (version 1.12). Data were normalized using global-scaling normalization (method: LogNormalize, scale.factor = 10.000). Highly variable genes (HVGs) were detected using *FindVariableGenes* function with default parameters. Then, normalized counts on HVGs were scaled and centered using *ScaleData* function with default parameters. Principal component analysis (PCA) was performed over the first ranked 2000 HVGs using *RunPCA* function (npcs = 50), and cluster detection was carried out with Louvain algorithm using 12 first PCA dimensions (decided by running *ElbowPlot*) at resolution = 0.2. Visualization and embedding were performed using UMAP over PCA using the 12 first PCA dimensions. Analysis was done first with young mice and after was re-analysed with both young and aged mice. Clusters with less than 1,000 nuclei in the sum of all conditions were annotated as other and excluded from further analysis. To annotated clusters to cell types, we used the single cell expression atlases of the mouse hippocampus and isocortex of Yao and colleagues (Yao et al., 2021). This was used as reference to annotate the study dataset by projecting their data structure to the query dataset using transfer anchors and UMAP projection via the *MapQuery* method within Seurat. We used robust cell type decomposition (RCTD)(*53*) to spatially annotate cell types in Slide-seqV2(*22*) data of the mouse hippocampus with annotations from our snRNA-seq, this correctly localized cell type predicted. These two approaches allowed us a robust and high-confidence calling of cell types. We used Seurat’s *FetchData* function to extract the normalized expression of some canonical IEGs (*Fos*, *Nr4a3*, *Egr2*, *Egr4*, *1700016P03Rik*, *Fosb*, *Arc*, *Egr1*, *Nr4a2*)(*54*) and divided cells in three activation states for each cluster: highly activated (IEGs expression sum > 2), lowly activated (IEGs expression sum < 2 & > 0) and not activated (IEGs expression sum = 0). Thus, when we filter activated cells, we choose highly and lowly activated cells. Moreover, *AddModuleScore* function from Seurat was used to identify IEGs signature. Plot1Cell(*55*) (version 0.0.0.9) facilitated the depiction of IEGs expression plots. Pseudobulk samples were generated for each cell type from each dataset using Seurat’s *AggregateExpression* function. Then, we performed differential expression analysis between experimental conditions for each cluster using DESeq2 (version 1.38.3). To determinate changes in cell composition between retrieval vs no retrieval and young vs aged, we used scProportionTest R package (version 0.0.0.9), which calculates the p-value of the magnitude difference of abundance for each cluster using a permutation test (number of permutations = 1000) and generates a confidence interval for the same using bootstrapping. To study the response of each cell type to memory retrieval Augur R package (version 1.0.3) was used, this compute “area under the receiver operating characteristic curve” (AUC) values. The neuron-neuron communication analysis between young and aged neuronal subtypes was performed using the R package NeuronChat(*25*) (version 1.0.0). We made a comparative analysis to confront the weights of the communication for individual interaction pairs and infer communication networks of relevant signalling pathways. Circle plot were plotted to show results. For gene ontology analysis, *Biological process*, *Cellular component* and *Molecular function* were performed using WebGestalt (WEB-based GEne SeT AnaLysis Toolkit(*51*); www.webgestalt.org), using over-representation analysis (ORA) as the enrichment method. Motif analysis of DEGs was performed using HOMER (version 4.11.1).

### Statistical analysis

All statistical analyses were conducted using GraphPad Prism (version 7.04, GraphPad Software, La Jolla CA) and RStudio (version 3.0.386). Statistical tests used in the study are indicated in the figure legend. When comparing two groups, Mann– Whitney U Statistic or two-way ANOVA was used. To determine differences between the median of a sample and a hypothesized median value, we used One-Sample Wilcoxon Test or one sample t-test. In all bar plots, the height represents the mean, and the error bars the standard error of mean (SEM).

## Data availability

The genomic data sets generated in this study can be accessed at the Gene Expression Omnibus (GEO) repository using the secure token <MVYJYSMEZVIZHGF> for dataset GSE289655.

## Acknowledgments

We thank Felix Leroy, Jose Vicente Sanchez-Mut and Rafael Alcala-Vida for critical reading of the manuscript. We thank the personnel of the Mouse facility, and the Omics core services at the Instituto de Neurociencias for their assistance. We also thank R. Olivares and C. Racovac for technical assistance. M.F-R. was recipient of a FPU fellowship from the Spanish Ministry of Education. M.A-N. was recipient of a fellowship from Generalitat Valenciana. A.B. research is supported by grants FCAIXA HR22-00394 from Fundación LaCaixa, PID2020-118169RB-I00 and PID2023-148442NB-I00 from AEI co-financed by ERDF, and CIPROM/2023/15 from the Generalitat Valenciana. The Instituto de Neurociencias is a “Centre of Excellence Severo Ochoa” (CEX2021-001165-S).

## Author contributions

Conceptualization, A.B.; Methodology, M.F-R., M.A-N., and F.M.; Software, M.F-R., A.V-P., and N.R.; Investigation, M.F-R., M.A-N., F.M. and J.F-A.; Data Curation and Visualization, M.F-R.; Writing - Original Draft, M.F-R. and A.B.; Supervision, N.R. and A.B.; Funding Acquisition, A.B.

## Declaration of generative AI and AI-assisted technologies in the writing process

During the preparation of this work the authors used ChatGTP to revise English grammar and usage. After using this tool/service, the authors reviewed and edited the content as needed and take full responsibility for the content of the publication.

## References

1. S. A. Josselyn, S. Tonegawa, Memory engrams: Recalling the past and imagining the future. Science 367 (2020).

2. A. Guskjolen, M. S. Cembrowski, Engram neurons: Encoding, consolidation, retrieval, and forgetting of memory. Mol. Psychiatry, doi: 10.1038/s41380-023-02137-5 (2023).

3. S. Tonegawa, M. D. Morrissey, T. Kitamura, The role of engram cells in the systems consolidation of memory. Nat. Rev. Neurosci. 19, 485–498 (2018).

4. M. Fuentes-Ramos, Á. Barco, “Unveiling Transcriptional and Epigenetic Mechanisms Within Engram Cells: Insights into Memory Formation and Stability” in Engrams: A Window into the Memory Trace, J. Gräff, S. Ramirez, Eds. (Springer International Publishing, Cham, 2024), pp. 111–129.

5. L. DeNardo, L. Luo, Genetic strategies to access activated neurons. Curr. Opin. Neurobiol. 45, 121–129 (2017).

6. C. J. Guenthner, K. Miyamichi, H. H. Yang, H. C. Heller, L. Luo, Permanent genetic access to transiently active neurons via TRAP: targeted recombination in active populations. Neuron 78, 773–784 (2013).

7. W. E. Allen, L. A. DeNardo, M. Z. Chen, C. D. Liu, K. M. Loh, L. E. Fenno, C. Ramakrishnan, K. Deisseroth, L. Luo, Thirst-associated preoptic neurons encode an aversive motivational drive. Science (80-. ). 357, 1149–1155 (2017).

8. L. A. DeNardo, C. D. Liu, W. E. Allen, E. L. Adams, D. Friedmann, L. Fu, C. J. Guenthner, M. Tessier-Lavigne, L. Luo, Temporal evolution of cortical ensembles promoting remote memory retrieval. Nat Neurosci 22, 460–469 (2019).

9. M. Fuentes-Ramos, M. Alaiz-Noya, A. Barco, Transcriptome and epigenome analysis of engram cells: Next-generation sequencing technologies in memory research. Neurosci. Biobehav. Rev. 127, 865–875 (2021).

10. N. Renier, Z. Wu, D. J. Simon, J. Yang, P. Ariel, M. Tessier-Lavigne, IDISCO: A simple, rapid method to immunolabel large tissue samples for volume imaging. Cell 159, 896– 910 (2014).

11. C. Kirst, S. Skriabine, A. Vieites-Prado, T. Topilko, P. Bertin, G. Gerschenfeld, F. Verny, P. Topilko, N. Michalski, M. Tessier-Lavigne, N. Renier, Mapping the Fine-Scale Organization and Plasticity of the Brain Vasculature. Cell 180, 780–795.e25 (2020).

12. N. Renier, E. L. Adams, C. Kirst, et al., Mapping of Brain Activity by Automated Volume Analysis of Immediate Early Genes. Cell 165, 1789–1802 (2016).

13. L. Madisen, T. A. Zwingman, S. M. Sunkin, S. W. Oh, H. A. Zariwala, H. Gu, L. L. Ng, R. D. Palmiter, M. J. Hawrylycz, A. R. Jones, E. S. Lein, H. Zeng, A robust and high-throughput Cre reporting and characterization system for the whole mouse brain. Nat. Neurosci. *2009 131* 13, 133–140 (2009).

14. J. K. Leutgeb, S. Leutgeb, M. B. Moser, E. I. Moser, Pattern separation in the dentate gyrus and CA3 of the hippocampus. Science (80-. ). 315, 961–966 (2007).

15. T. Kitamura, S. K. Ogawa, D. S. Roy, T. Okuyama, M. D. Morrissey, L. M. Smith, R. L. Redondo, S. Tonegawa, Engrams and circuits crucial for systems consolidation of a memory. Science (80-. ). 356, 73–78 (2017).

16. J. H. Lee, W. Bin Kim, E. H. Park, J. H. Cho, Neocortical synaptic engrams for remote contextual memories. Nat. Neurosci. *2022 262* 26, 259–273 (2022).

17. R. J. Robitsek, N. J. Fortin, T. K. Ming, M. Gallagher, H. Eichenbaum, Cognitive Aging: A Common Decline of Episodic Recollection and Spatial Memory in Rats. J. Neurosci. 28, 8945–8954 (2008).

18. M. Gallagher, P. R. Rapp, The use of animal models to study the effects of aging on cognition. Annu. Rev. Psychol. 48, 339–370 (1997).

19. Z. Yao, C. T. J. van Velthoven, T. N. Nguyen, et al., A taxonomy of transcriptomic cell types across the isocortex and hippocampal formation. Cell 184, 3222–3241.e26 (2021).

20. M. B. Chen, X. Jiang, S. R. Quake, T. C. Südhof, Persistent transcriptional programmes are associated with remote memory. Nature 587, 437–442 (2020).

21. W. Sun, Z. Liu, X. Jiang, M. B. Chen, H. Dong, J. Liu, T. C. Südhof, S. R. Quake, Spatial transcriptomics reveal neuron–astrocyte synergy in long-term memory. Nat. *2024 6278003* 627, 374–381 (2024).

22. R. R. Stickels, E. Murray, P. Kumar, J. Li, J. L. Marshall, D. J. Di Bella, P. Arlotta, E. Z. Macosko, F. Chen, Highly sensitive spatial transcriptomics at near-cellular resolution with Slide-seqV2. Nat. Biotechnol. 39, 313–319 (2021).

23. G. Miyoshi, A. Young, T. Petros, et al., Prox1 Regulates the Subtype-Specific Development of Caudal Ganglionic Eminence-Derived GABAergic Cortical Interneurons. J. Neurosci. 35, 12869–12889 (2015).

24. J. W. Squair, M. A. Skinnider, M. Gautier, L. J. Foster, G. Courtine, Prioritization of cell types responsive to biological perturbations in single-cell data with Augur. Nat. Protoc. *2021 168* 16, 3836–3873 (2021).

25. W. Zhao, K. G. Johnston, H. Ren, X. Xu, Q. Nie, Inferring neuron-neuron communications from single-cell transcriptomics through NeuronChat. Nat. Commun. *2023 141* 14, 1–16 (2023).

26. P. Rao-Ruiz, J. J. Couey, I. M. Marcelo, C. G. Bouwkamp, D. E. Slump, M. R. Matos, R. J. van der Loo, G. J. Martins, M. van den Hout, W. F. van IJcken, R. M. Costa, M. C. van den Oever, S. A. Kushner, Engram-specific transcriptome profiling of contextual memory consolidation. Nat. Commun. 10, 2232 (2019).

27. Y. Gao, M. Shen, J. C. Gonzalez, Q. Dong, S. Kannan, J. T. Hoang, B. E. Eisinger, J. Pandey, S. Javadi, Q. Chang, D. Wang, L. Overstreet-Wadiche, X. Zhao, RGS6 mediates effects of voluntary running on adult hippocampal neurogenesis. Cell Rep. 32, 107997 (2020).

28. A. Marco, H. S. Meharena, V. Dileep, R. M. Raju, J. Davila-Velderrain, A. L. Zhang, C. Adaikkan, J. Z. Young, F. Gao, M. Kellis, L.-H. Tsai, Mapping the epigenomic and transcriptomic interplay during memory formation and recall in the hippocampal engram ensemble. Nat. Neurosci. 23, 1606–1617 (2020).

29. N. Chiaruttini, C. Castoldi, L. M. Requie, C. Camarena-Delgado, B. dal Bianco, J. Gräff, A. Seitz, B. A. Silva, ABBA, a novel tool for whole-brain mapping, reveals brain-wide differences in immediate early genes induction following learning. bioRxiv, 2024.09.06.611625 (2024).

30. T. Stefanelli, C. Bertollini, C. Lüscher, D. Muller, P. Mendez, Hippocampal Somatostatin Interneurons Control the Size of Neuronal Memory Ensembles. Neuron 89, 1074–1085 (2016).

31. G. Vetere, J. W. Kenney, L. M. Tran, F. Xia, P. E. Steadman, J. Parkinson, S. A. Josselyn, P. W. Frankland, Chemogenetic Interrogation of a Brain-wide Fear Memory Network in Mice. Neuron 94, 363–374.e4 (2017).

32. D. S. Roy, Y.-G. G. Park, M. E. Kim, et al., Brain-wide mapping reveals that engrams for a single memory are distributed across multiple brain regions. Nat. Commun. 13, 1799 (2022).

33. B. N. Jaeger, S. B. Linker, S. L. Parylak, J. J. Barron, I. S. Gallina, C. D. Saavedra, C. Fitzpatrick, C. K. Lim, S. T. Schafer, B. Lacar, S. Jessberger, F. H. Gage, A novel environment-evoked transcriptional signature predicts reactivity in single dentate granule neurons. Nat. Commun. 9, 3084 (2018).

34. J. Lisman, K. Cooper, M. Sehgal, A. J. Silva, Memory formation depends on both synapse-specific modifications of synaptic strength and cell-specific increases in excitability. Nat. Neurosci. *2018 213* 21, 309–314 (2018).

35. G. Vetere, L. Restivo, C. J. Cole, P. J. Ross, M. Ammassari-Teule, S. A. Josselyn, P. W. Frankland, Spine growth in the anterior cingulate cortex is necessary for the consolidation of contextual fear memory. Proc. Natl. Acad. Sci. U. S. A., doi: 10.1073/pnas.1016275108 (2011).

36. E. Benito, A. Barco, CREB’s control of intrinsic and synaptic plasticity: implications for CREB-dependent memory models. Trends Neurosci. 33, 230–240 (2010).

37. X. Sun, M. J. Bernstein, M. Meng, S. Rao, A. T. Sørensen, L. Yao, X. Zhang, P. O. Anikeeva, Y. Lin, Functionally Distinct Neuronal Ensembles within the Memory Engram. Cell 181, 410–423.e17 (2020).

38. A. Besnard, B. Galan-Rodriguez, P. Vanhoutte, J. Caboche, Elk-1 a transcription factor with multiple facets in the brain. Front. Neurosci. 5, 9281 (2011).

39. W. E. Allen, T. R. Blosser, Z. A. Sullivan, C. Dulac, X. Zhuang, Molecular and spatial signatures of mouse brain aging at single-cell resolution. Cell 186, 194–208.e18 (2023).

40. K. Jin, Z. Yao, C. T. J. van Velthoven, et al., Brain-wide cell-type-specific transcriptomic signatures of healthy ageing in mice. Nat. 2025, 1–15 (2025).

41. A. Sziraki, Z. Lu, J. Lee, et al., A global view of aging and Alzheimer’s pathogenesis-associated cell population dynamics and molecular signatures in human and mouse brains. Nat. Genet. 2023 5512 55, 2104–2116 (2023).

42. M. Ximerakis, S. L. Lipnick, B. T. Innes, et al., Single-cell transcriptomic profiling of the aging mouse brain. Nat. Neurosci. 2019 2210 22, 1696–1708 (2019).

43. S. Maren, K. L. Phan, I. Liberzon, The contextual brain: implications for fear conditioning, extinction and psychopathology. Nat. Rev. Neurosci. *2013 146* 14, 417–428 (2013).

44. T. D. Goode, K. Z. Tanaka, A. Sahay, T. J. McHugh, An Integrated Index: Engrams, Place Cells, and Hippocampal Memory. Neuron 107, 805–820 (2020).

45. G. Erdmann, G. Schütz, S. Berger, Inducible gene inactivation in neurons of the adult mouse forebrain. BMC Neurosci. 8, 63 (2007).

46. A. Mo, E. A. Mukamel, F. P. Davis, C. Luo, G. L. Henry, S. Picard, M. A. Urich, J. R. Nery, T. J. Sejnowski, R. Lister, S. R. Eddy, J. R. Ecker, J. Nathans, Epigenomic Signatures of Neuronal Diversity in the Mammalian Brain. Neuron 86, 1369–1384 (2015).

47. A. Fiorenza, J. P. Lopez-Atalaya, V. Rovira, M. Scandaglia, E. Geijo-Barrientos, A. Barco, Blocking miRNA Biogenesis in Adult Forebrain Neurons Enhances Seizure Susceptibility, Fear Memory, and Food Intake by Increasing Neuronal Responsiveness. Cereb. Cortex 26, 1619–1633 (2016).

48. W. Bin Kim, J. H. Cho, Encoding of contextual fear memory in hippocampal–amygdala circuit. Nat. Commun. *2020 111* 11, 1–22 (2020).

49. T. Smith, A. Heger, I. Sudbery, UMI-tools: modeling sequencing errors in Unique Molecular Identifiers to improve quantification accuracy. Genome Res. 27, 491–499 (2017).

50. A. Dobin, C. A. Davis, F. Schlesinger, J. Drenkow, C. Zaleski, S. Jha, P. Batut, M. Chaisson, T. R. Gingeras, STAR: ultrafast universal RNA-seq aligner. Bioinformatics 29, 15–21 (2013).

51. Y. Liao, J. Wang, E. J. Jaehnig, Z. Shi, B. Zhang, WebGestalt 2019: gene set analysis toolkit with revamped UIs and APIs. Nucleic Acids Res. 47, W199–W205 (2019).

52. F. Koopmans, P. van Nierop, M. Andres-Alonso, et al., SynGO: An Evidence-Based, Expert-Curated Knowledge Base for the Synapse. Neuron 103, 217–234.e4 (2019).

53. D. M. Cable, E. Murray, L. S. Zou, A. Goeva, E. Z. Macosko, F. Chen, R. A. Irizarry, Robust decomposition of cell type mixtures in spatial transcriptomics. Nat. Biotechnol. 40, 517–526 (2022).

54. J. Fernandez-Albert, M. Lipinski, M. T. Lopez-Cascales, M. J. Rowley, A. M. Martin-Gonzalez, B. del Blanco, V. G. Corces, A. Barco, Immediate and deferred epigenomic signatures of in vivo neuronal activation in mouse hippocampus. Nat Neurosci 22, 1718–1730 (2019).

55. H. Wu, R. G. Villalobos, X. Yao, D. Reilly, T. Chen, M. Rankin, E. Myshkin, M. D. Breyer, A. D. Humphreys, Mapping the single-cell transcriptomic response of murine diabetic kidney disease to therapies. Cell Metab. 34, 1064–1078.e6 (2022).

