## Supplementary material for "Aging alters the distribution, stability, and transcriptional signature of engram cells": Supp Material and Figures

• Supplementary Figures S1-S15 (pages 2-31)

• Supplementary Files:

**Table S1. ClearMap2 data comparing SUN1-GFP levels of FT and HC conditions in 378 brain regions.** Each region is sorted by p values (ascending order). Mean, SD, p values and log2FoldChange (FC).

**Table S2. ClearMap2 data comparing FOS levels of A-A and A-B conditions in 378 brain regions.** Each region is sorted by p values (ascending order). Mean, SD, p values and log2FC.

**Table S3. ClearMap2 data comparing FOS levels of A-1-A and A-3-A conditions in 378 brain regions.** Each region is sorted by p values (ascending order). Mean, SD, p values and log2FC.

**Table S4. ClearMap2 data comparing FOS levels of A-A and A-B conditions in aged mice for 378 brain regions.** Each region is sorted by p values (ascending order). Mean, SD, p-value and log2FC.

**Table S5. snRNA-seq and nuRNA-seq samples and experimental metadata.** 2 sheets: (i) snRNA-seq sample description; (ii) nuRNA-seq sample description.

**Table S6. Result of differential expression analyses for DEGs in R vs NR comparison for young mice.** 10 sheets, one for each cell type, presenting gene symbol, log2FC and p-adjusted value.

**Table S7. List of DEGs in the comparison R vs NR in young mice, grouped into three clusters:** UIG, Repressed genes and Memory-induced excitatory-specific genes.

**Table S8. Result of differential expression analyses for all genes in R vs NR comparison for young and aged mice.** 10 sheets, one for each cell type, presenting gene symbol, log2FC and p-adjusted value.

**Table S9. Differentially expressed genes in nuRNA-seq.** 4 sheets: (i) DEGs that are upregulated in reactivated cells and their assignment to clusters; (ii) DEGs in the comparison of Reactivated vs 1$^{st}$ experience cells; (iii) DEGs in the comparison of 2$^{nd}$ experience vs reactivated cells; and (iv) DEGs in the comparison of 2$^{nd}$ experience vs 1$^{st}$ experience cells.

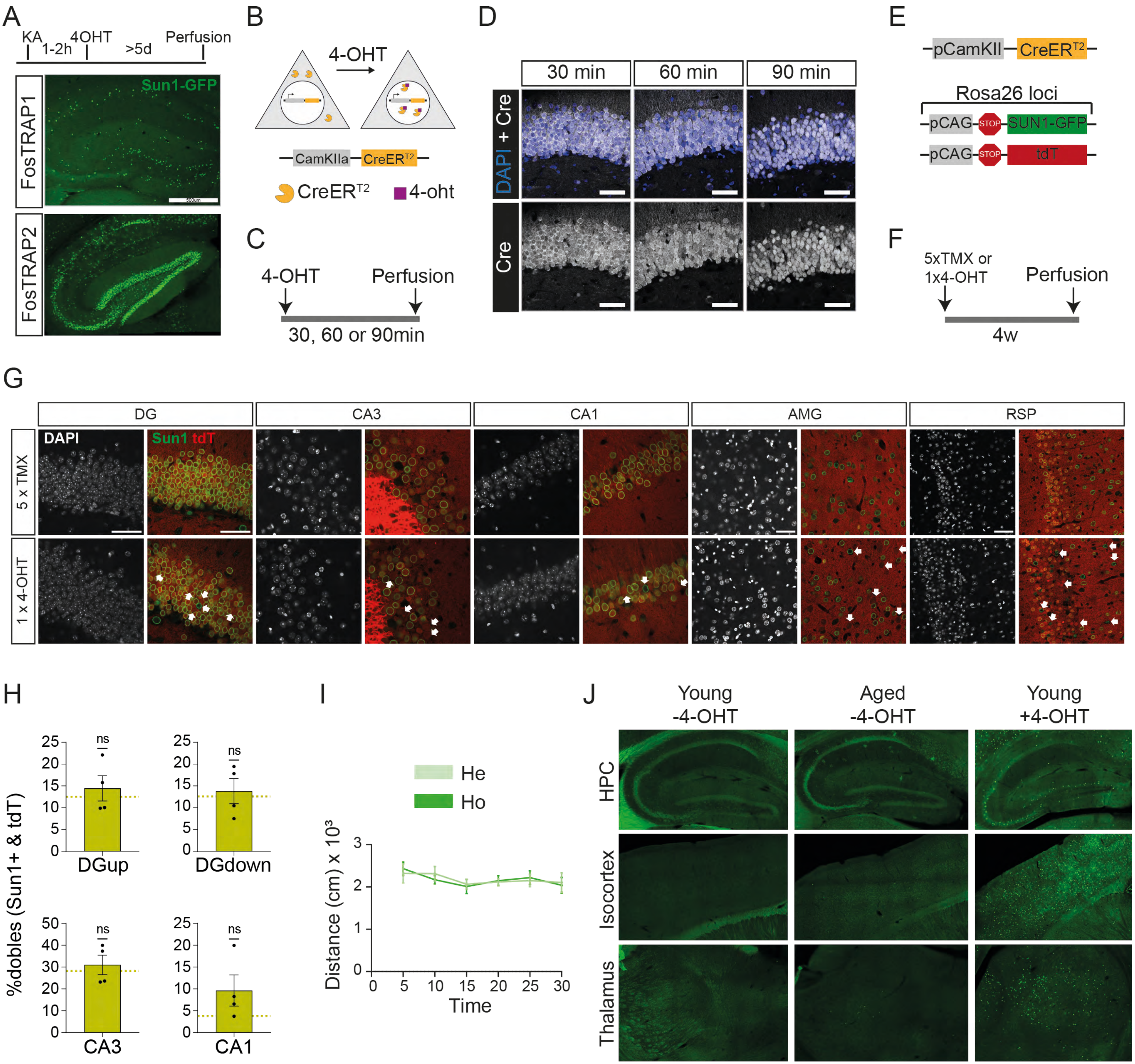
A
KA
1-2h
4OHT
>5d
Perfusion
Sun1-GFP
FosTRAP1
FosTRAP2
B
4-OHT
CamKIIa
CreER$^{T2}$
4-oht
C
4-OHT
Perfusion
30, 60 or 90min
D
30 min
60 min
90 min
DAPI + Cre
Cre
E
pCamKII
CreER$^{T2}$
Rosa26 loci
pCAG
STOP
SUN1-GFP
tdT
F
5xTMX or
1x4-OHT
Perfusion
4w
G
DG
CA3
CA1
AMG
RSP
DAPI
Sun1 tdT
5 x TMX
1 x 4-OHT
H
%dobles (Sun1+ & tdT)
ns
DGup
DGdown
CA3
CA1
I
He
Ho
Distance (cm) x 10$^3$
Time
J
Young -4-OHT
Aged -4-OHT
Young +4-OHT
HPC
Isocortex
Thalamus

**Supplementary Figure S1 (related to Figure 1). Characterization of the TRAP2 reporter strain. A.** FosTRAP1 and FosTRAP2 mice carry a Cre-dependent SUN1-GFP reporter in one Rosa26 allele, enabling nuclear envelope labeling in FOS-expressing cells upon 4-OHT administration. KA-induced neuronal activation labeled with 4-OHT reveals higher sensitivity in the FosTRAP2 line. Scale bar: 500 µm. **B.** In the tamoxifen (TAM)-inducible Cre-driver line CaMKII-CreERT2, 4-OHT treatment enables nuclear translocation of Cre recombinase. **C.** Experimental design. **D.** Confocal images of dentate gyrus granule cells expressing Cre recombinase. **E.** CaMKII-CreERT2 mice were crossed with mice carrying SUN1-GFP in a Rosa26 allele and tdTomato in the other. **F.** Experimental design to test TAM/4-OHT dosage as a limiting factor for reliable labeling active neurons. **G.** Confocal images showing reporter expression in various hippocampal regions, the amygdala (AMG), and retrosplenial cortex (RSP) after different 4-OHT doses. Scale bar: 50 µm. **H.** Quantification of double labeling with SUN1-GFP and tdTomato in hippocampal layers compared to chance levels (One-sample Wilcoxon test). **I.** Distance traveled during the first 30 minutes of novel environment exposure (two-way ANOVA: not significant, $P > 0.05$). **J.** Confocal images from young and aged TRAP2 mice without 4-OHT injection, and from young TRAP2 mice with 4-OHT.

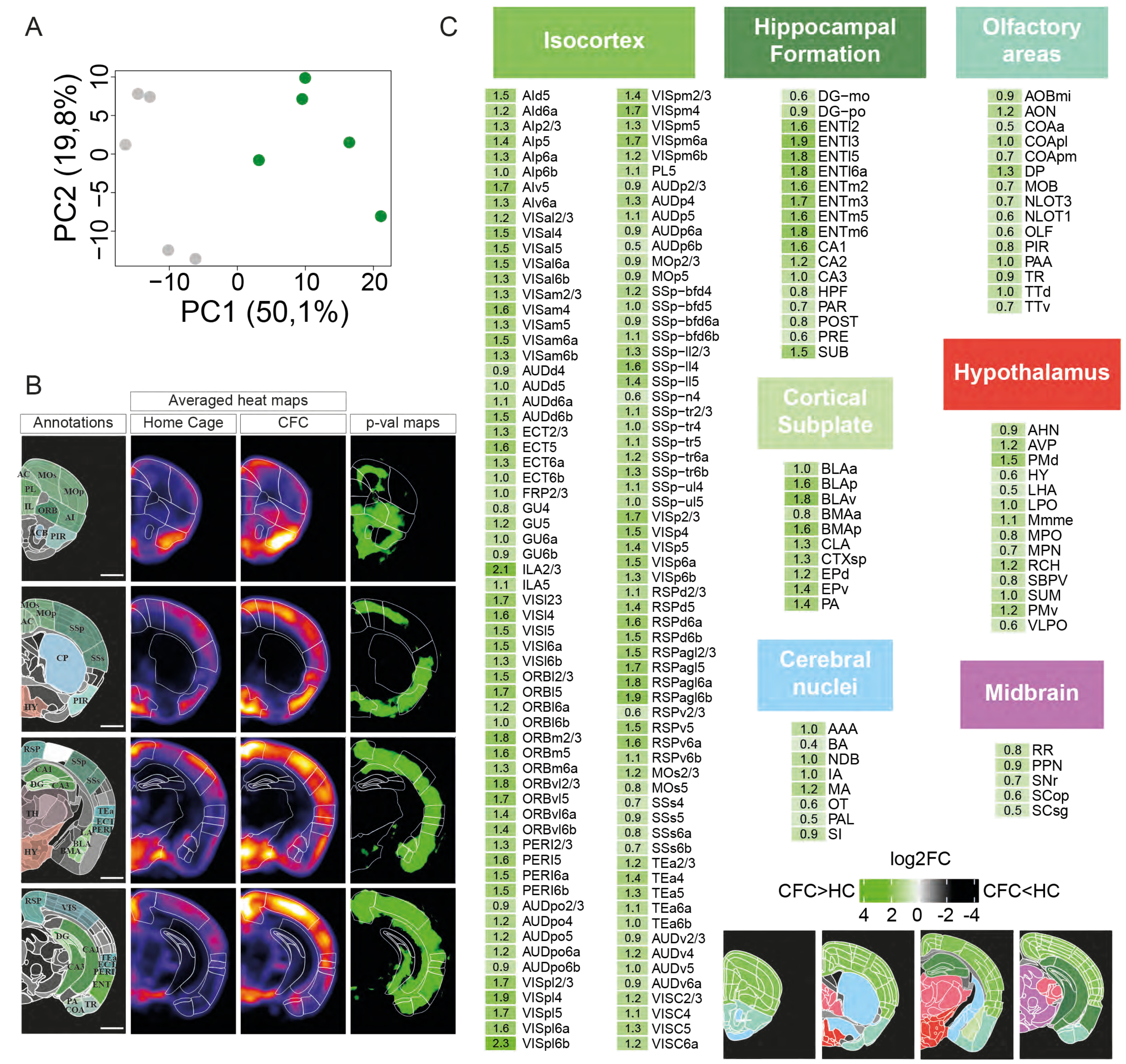
A
PC2 (19,8%)
PC1 (50,1%)
−10 −5 0 5 10
−10 0 10 20
B
Averaged heat maps
Annotations
Home Cage
CFC
p-val maps
C
Isocortex
1.5 AId5
1.2 AId6a
1.3 AIp2/3
1.4 AIp5
1.3 AIp6a
1.0 AIp6b
1.7 AIv5
1.3 AIv6a
1.2 VISal2/3
1.5 VISal4
1.5 VISal5
1.5 VISal6a
1.3 VISal6b
1.3 VISam2/3
1.6 VISam4
1.3 VISam5
1.5 VISam6a
1.3 VISam6b
0.9 AUDd4
1.0 AUDd5
1.1 AUDd6a
1.5 AUDd6b
1.3 ECT2/3
1.6 ECT5
1.3 ECT6a
1.0 ECT6b
1.0 FRP2/3
0.8 GU4
1.2 GU5
1.0 GU6a
0.9 GU6b
2.1 ILA2/3
1.1 ILA5
1.7 VISl23
1.6 VISl4
1.5 VISl5
1.5 VISl6a
1.3 VISl6b
1.5 ORBl2/3
1.7 ORBl5
1.2 ORBl6a
1.0 ORBl6b
1.8 ORBm2/3
1.6 ORBm5
1.3 ORBm6a
1.8 ORBvl2/3
1.7 ORBvl5
1.4 ORBvl6a
1.4 ORBvl6b
1.3 PERI2/3
1.6 PERI5
1.5 PERI6a
1.5 PERI6b
0.9 AUDpo2/3
1.2 AUDpo4
1.2 AUDpo5
1.2 AUDpo6a
0.9 AUDpo6b
1.7 VISpl2/3
1.9 VISpl4
1.7 VISpl5
1.6 VISpl6a
2.3 VISpl6b
1.4 VISpm2/3
1.7 VISpm4
1.3 VISpm5
1.7 VISpm6a
1.2 VISpm6b
1.1 PL5
0.9 AUDp2/3
1.3 AUDp4
1.1 AUDp5
0.9 AUDp6a
0.5 AUDp6b
0.9 MOp2/3
0.9 MOp5
1.2 SSp−bfd4
1.0 SSp−bfd5
0.9 SSp−bfd6a
1.1 SSp−bfd6b
1.3 SSp−ll2/3
1.6 SSp−ll4
1.4 SSp−ll5
0.6 SSp−n4
1.1 SSp−tr2/3
1.0 SSp−tr4
1.1 SSp−tr5
1.2 SSp−tr6a
1.3 SSp−tr6b
1.1 SSp−ul4
1.0 SSp−ul5
1.7 VISp2/3
1.5 VISp4
1.4 VISp5
1.5 VISp6a
1.3 VISp6b
1.1 RSPd2/3
1.4 RSPd5
1.6 RSPd6a
1.5 RSPd6b
1.5 RSPagl2/3
1.7 RSPagl5
1.8 RSPagl6a
1.9 RSPagl6b
0.6 RSPv2/3
1.5 RSPv5
1.6 RSPv6a
1.1 RSPv6b
1.2 MOs2/3
0.8 MOs5
0.7 SSs4
0.9 SSs5
0.8 SSs6a
0.7 SSs6b
1.2 TEa2/3
1.4 TEa4
1.3 TEa5
1.1 TEa6a
1.0 TEa6b
0.9 AUDv2/3
1.2 AUDv4
1.0 AUDv5
0.9 AUDv6a
1.2 VISC2/3
1.1 VISC4
1.3 VISC5
1.2 VISC6a
Hippocampal Formation
0.6 DG−mo
0.9 DG−po
1.6 ENTl2
1.9 ENTl3
1.8 ENTl5
1.8 ENTl6a
1.6 ENTm2
1.7 ENTm3
1.6 ENTm5
1.8 ENTm6
1.6 CA1
1.2 CA2
1.0 CA3
0.8 HPF
0.7 PAR
0.8 POST
0.6 PRE
1.5 SUB
Cortical Subplate
1.0 BLAa
1.6 BLAp
1.8 BLAv
0.8 BMAa
1.6 BMAp
1.3 CLA
1.3 CTXsp
1.2 EPd
1.4 EPv
1.4 PA
Cerebral nuclei
1.0 AAA
0.4 BA
1.0 NDB
1.0 IA
1.2 MA
0.6 OT
0.5 PAL
0.9 SI
Olfactory areas
0.9 AOBmi
1.2 AON
0.5 COAa
1.0 COApl
0.7 COApm
1.3 DP
0.7 MOB
0.7 NLOT3
0.6 NLOT1
0.6 OLF
0.8 PIR
1.0 PAA
0.9 TR
1.0 TTd
0.7 TTv
Hypothalamus
0.9 AHN
1.2 AVP
1.5 PMd
0.6 HY
0.5 LHA
1.0 LPO
1.1 Mmme
0.8 MPO
0.7 MPN
1.2 RCH
0.8 SBPV
1.0 SUM
1.2 PMv
0.6 VLPO
Midbrain
0.8 RR
0.9 PPN
0.7 SNr
0.6 SCop
0.5 SCsg
log2FC
CFC>HC
CFC<HC
4 2 0 -2 -4

**Supplementary Figure S2 (related to Figure 2). Brain subregions with cells tagged during contextual fear conditioning (CFC). A.** Principal component analysis (PCA) of SUN1-GFP expression across individual mice. The CFC group (green) separates from the home cage (HC) group (gray), indicating a distinct activation pattern. **B.** Coronal sections from the automated analysis of SUN1-GFP–positive cell distribution. Shown are the reference atlas annotations, averaged density maps per condition (5 brains per group), and p-value maps highlighting regions with significantly increased SUN1-GFP labeling in the CFC group (green). Scale bars: 500 µm. **C.** A total of 197 brain subregions exhibited significantly higher SUN1-GFP–positive cell counts in the CFC group compared to controls.

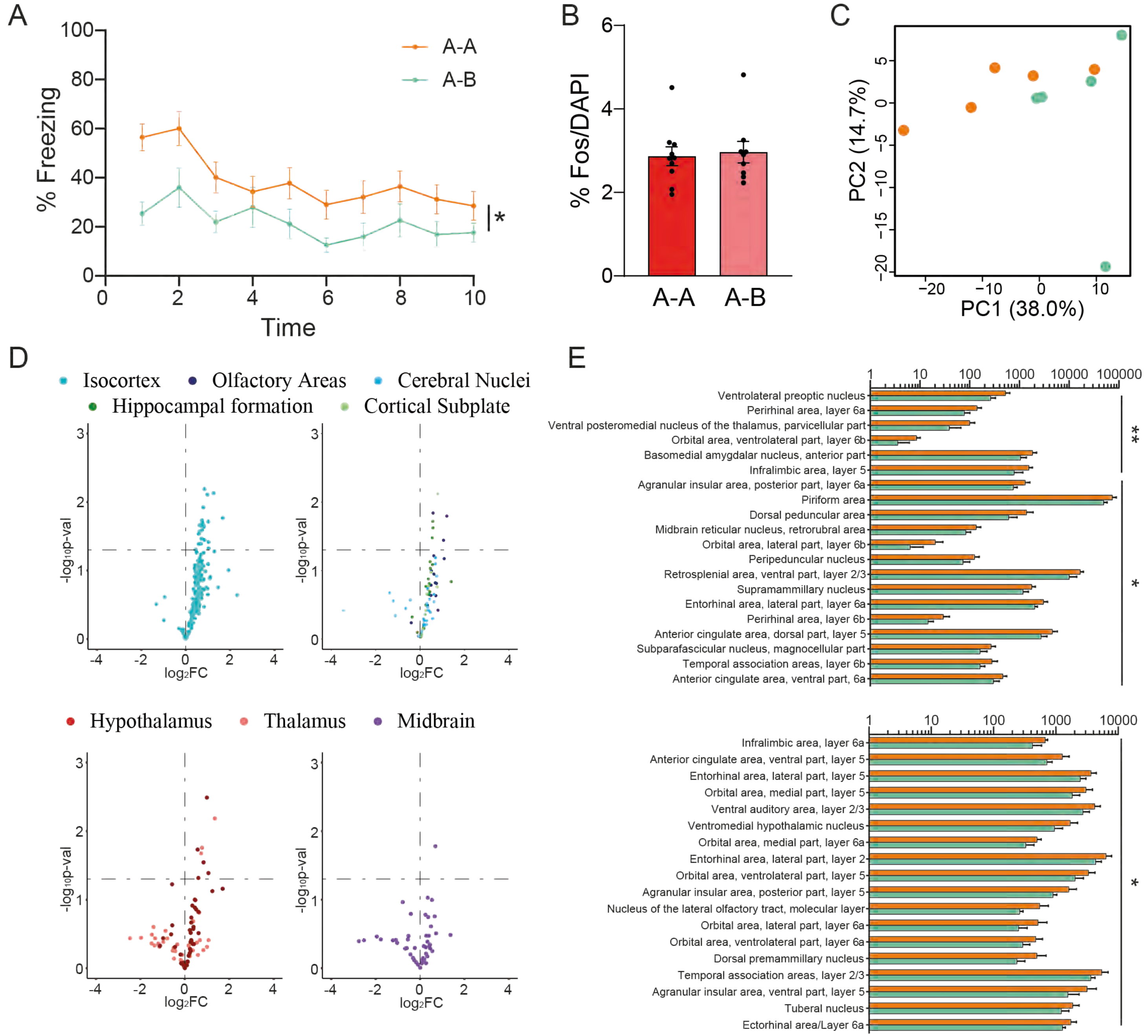
A
% Freezing
Time
A-A
A-B
*
B
% Fos/DAPI
A-A
A-B
C
PC1 (38.0%)
PC2 (14.7%)
D
Isocortex
Olfactory Areas
Cerebral Nuclei
Hippocampal formation
Cortical Subplate
Hypothalamus
Thalamus
Midbrain
-log10p-val
log2FC
E
Ventrolateral preoptic nucleus
Perirhinal area, layer 6a
Ventral posteromedial nucleus of the thalamus, parvicellular part
Orbital area, ventrolateral part, layer 6b
Basomedial amygdalar nucleus, anterior part
Infralimbic area, layer 5
Agranular insular area, posterior part, layer 6a
Piriform area
Dorsal peduncular area
Midbrain reticular nucleus, retrorubral area
Orbital area, lateral part, layer 6b
Peripeduncular nucleus
Retrosplenial area, ventral part, layer 2/3
Supramammillary nucleus
Entorhinal area, lateral part, layer 6a
Perirhinal area, layer 6b
Anterior cingulate area, dorsal part, layer 5
Subparafascicular nucleus, magnocellular part
Temporal association areas, layer 6b
Anterior cingulate area, ventral part, 6a
**
*
Infralimbic area, layer 6a
Anterior cingulate area, ventral part, layer 5
Entorhinal area, lateral part, layer 5
Orbital area, medial part, layer 5
Ventral auditory area, layer 2/3
Ventromedial hypothalamic nucleus
Orbital area, medial part, layer 6a
Entorhinal area, lateral part, layer 2
Orbital area, ventrolateral part, layer 5
Agranular insular area, posterior part, layer 5
Nucleus of the lateral olfactory tract, molecular layer
Orbital area, lateral part, layer 6a
Orbital area, ventrolateral part, layer 6a
Dorsal premammillary nucleus
Temporal association areas, layer 2/3
Agranular insular area, ventral part, layer 5
Tuberal nucleus
Ectorhinal area/Layer 6a
*

**Supplementary Figure S3 (related to Figure 2). Memory recall induces greater FOS expression than novelty exploration. A.** Freezing levels during memory retrieval measured in 1-minute intervals over a 10-minute period (two-way ANOVA: *$P_{condition}$ < 0.05). **B.** Percentage of $FOS^{+}$ cells relative to total number of $DAPI^{+}$ cells in DG (A-A: n=10; A-B: n=9; Mann–Whitney U Statistic). **C.** PCA of FOS expression across individual mice. **D.** Volcano plots showing differential FOS expression across brain regions. **E.** Automated quantification of $FOS^{+}$ cell counts (mean ± SEM) by anatomical region, sorted by *p*-value (*n* = 5 per condition). **P* < 0.05, ** *P* < 0.01, *** *P* < 0.001.

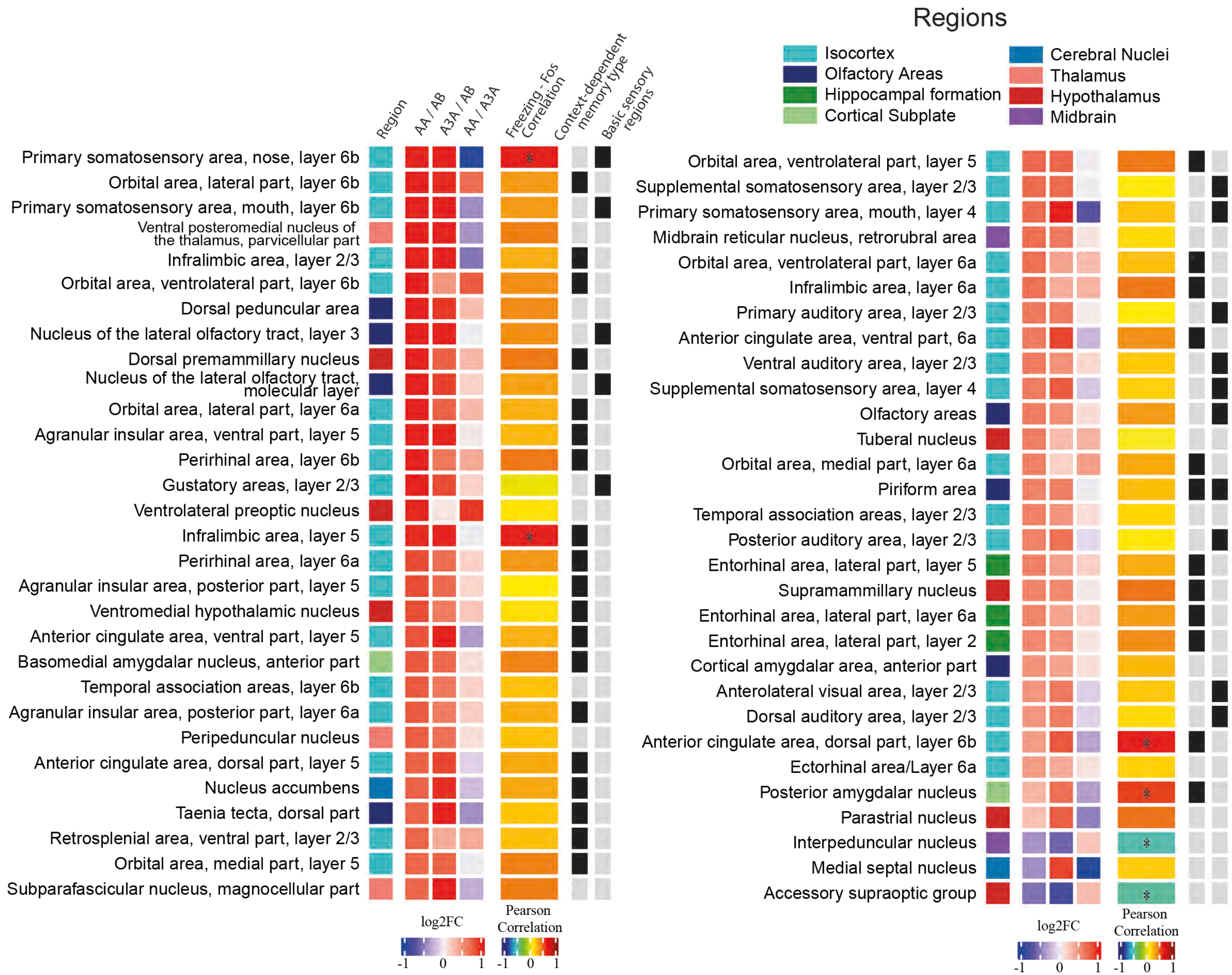

Regions
Isocortex
Olfactory Areas
Hippocampal formation
Cortical Subplate
Cerebral Nuclei
Thalamus
Hypothalamus
Midbrain
Region
AA / AB
A3A / AB
AA / A3A
Freezing - Fos Correlation
Context-dependent memory type
Basic sensory regions
Primary somatosensory area, nose, layer 6b
Orbital area, lateral part, layer 6b
Primary somatosensory area, mouth, layer 6b
Ventral posteromedial nucleus of the thalamus, parvicellular part
Infralimbic area, layer 2/3
Orbital area, ventrolateral part, layer 6b
Dorsal peduncular area
Nucleus of the lateral olfactory tract, layer 3
Dorsal premammillary nucleus
Nucleus of the lateral olfactory tract, molecular layer
Orbital area, lateral part, layer 6a
Agranular insular area, ventral part, layer 5
Perirhinal area, layer 6b
Gustatory areas, layer 2/3
Ventrolateral preoptic nucleus
Infralimbic area, layer 5
Perirhinal area, layer 6a
Agranular insular area, posterior part, layer 5
Ventromedial hypothalamic nucleus
Anterior cingulate area, ventral part, layer 5
Basomedial amygdalar nucleus, anterior part
Temporal association areas, layer 6b
Agranular insular area, posterior part, layer 6a
Peripeduncular nucleus
Anterior cingulate area, dorsal part, layer 5
Nucleus accumbens
Taenia tecta, dorsal part
Retrosplenial area, ventral part, layer 2/3
Orbital area, medial part, layer 5
Subparafascicular nucleus, magnocellular part
log2FC
-1 0 1
Pearson Correlation
-1 0 1
Orbital area, ventrolateral part, layer 5
Supplemental somatosensory area, layer 2/3
Primary somatosensory area, mouth, layer 4
Midbrain reticular nucleus, retrorubral area
Orbital area, ventrolateral part, layer 6a
Infralimbic area, layer 6a
Primary auditory area, layer 2/3
Anterior cingulate area, ventral part, 6a
Ventral auditory area, layer 2/3
Supplemental somatosensory area, layer 4
Olfactory areas
Tuberal nucleus
Orbital area, medial part, layer 6a
Piriform area
Temporal association areas, layer 2/3
Posterior auditory area, layer 2/3
Entorhinal area, lateral part, layer 5
Supramammillary nucleus
Entorhinal area, lateral part, layer 6a
Entorhinal area, lateral part, layer 2
Cortical amygdalar area, anterior part
Anterolateral visual area, layer 2/3
Dorsal auditory area, layer 2/3
Anterior cingulate area, dorsal part, layer 6b
Ectorhinal area/Layer 6a
Posterior amygdalar nucleus
Parastrial nucleus
Interpeduncular nucleus
Medial septal nucleus
Accessory supraoptic group
log2FC
-1 0 1
Pearson Correlation
-1 0 1

**Supplementary Figure S4 (related to Figure 2). Brain subregions with differential FOS expression during fear recall or context discrimination.** Sixty brain subregions are shown with their log2FC across three experimental comparisons. The plot also displays correlations between FOS levels and freezing behavior (*P* < 0.05), and identifies subregions previously associated with contextual memory or basic sensory areas (both labelled in black).

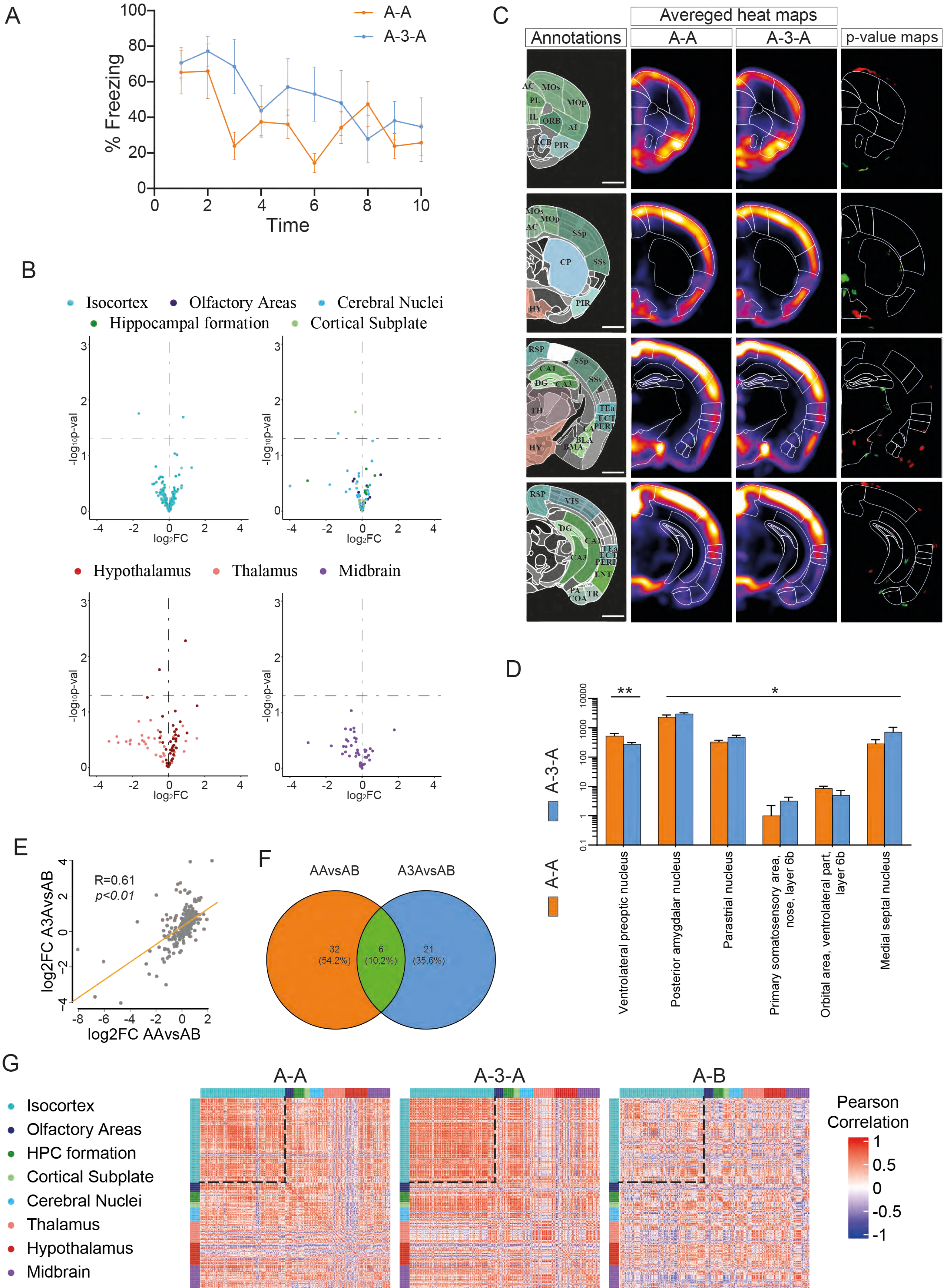
A
A-A
A-3-A
% Freezing
Time
C
Avereged heat maps
Annotations
A-A
A-3-A
p-value maps
B
Isocortex
Olfactory Areas
Cerebral Nuclei
Hippocampal formation
Cortical Subplate
-log10p-val
log2FC
Hypothalamus
Thalamus
Midbrain
D
A-3-A
A-A
Ventrolateral preoptic nucleus
Posterior amygdalar nucleus
Parastrial nucleus
Primary somatosensory area, nose, layer 6b
Orbital area, ventrolateral part, layer 6b
Medial septal nucleus
E
R=0.61
p<0.01
log2FC A3AvsAB
log2FC AAvsAB
F
AAvsAB
A3AvsAB
32 (54.2%)
6 (10.2%)
21 (35.6%)
G
A-A
A-3-A
A-B
Isocortex
Olfactory Areas
HPC formation
Cortical Subplate
Cerebral Nuclei
Thalamus
Hypothalamus
Midbrain
Pearson Correlation

**Supplementary Figure S5 (related to Figure 2). Cellular activation remains stable over time but exhibits changes in inter-regional coupling patterns. A.** Freezing behavior during a 10-minute retrieval session was similar between recent memory (A–A) and remote memory (A–3w–A) groups (two-way ANOVA: not significant, $P_{condition} > 0.05$). **B.** Volcano plots comparing FOS expression between remote (A–3w–A) and recent (A–A) memory retrieval. **C.** Coronal views of automated FOS$^{+}$ cell mapping. Panels show atlas-based annotations, averaged density maps (5 brains per group), and p-value maps indicating significantly increased FOS expression in A–A (red) and A–3w–A (green). **D.** Automated quantification of FOS+ cell counts (mean ± SEM) across anatomical regions, sorted by *p*-value. **E.** Log2FC in FOS expression for each brain subregion (gray dots), comparing fear retrieval versus novel context exploration: A–A vs. A–B (x-axis) and A–3w–A vs. A–B (y-axis). **F.** Overlap of subregions showing increased activation in both the A–3w–A vs. A–B and A–A vs. A–B comparisons. **G.** Subregion-by-subregion correlation matrices of FOS expression across brain regions for each experimental group. Activity levels were normalized by regional volume.

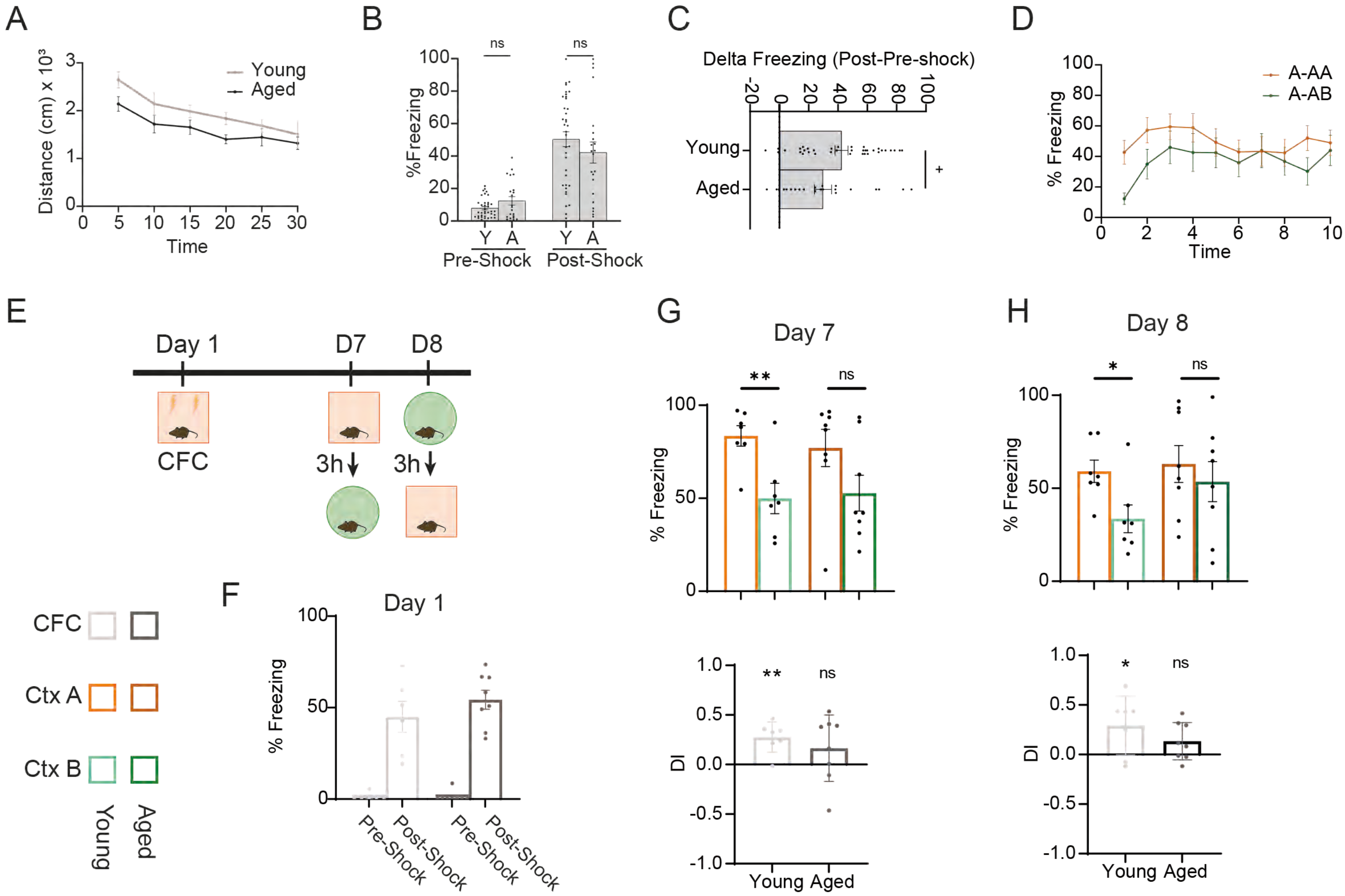
A
Young
Aged
Distance (cm) x 10³
Time
B
ns
ns
%Freezing
Y A Y A
Pre-Shock Post-Shock
C
Delta Freezing (Post-Pre-shock)
Young
Aged
+
D
A-AA
A-AB
% Freezing
Time
E
Day 1
D7
D8
CFC
3h
3h
CFC
Ctx A
Ctx B
Young
Aged
F
Day 1
% Freezing
Pre-Shock
Post-Shock
Pre-Shock
Post-Shock
G
Day 7
**
ns
% Freezing
DI
**
ns
Young Aged
H
Day 8
*
ns
% Freezing
DI
*
ns
Young Aged

**Supplementary Figure S6 (related to Figure 3). Contextual discrimination and cognitive flexibility are affected in aged mice. A.** Distance travelled during the first 30 minutes of the novel exposure (two-way ANOVA: ns $P_{condition} > 0.05$). **B.** Differences in the freezing response of young (Y) and aged (A) mice during pre- and post-shock (Mann–Whitney U Statistic). **C.** Differences in delta freezing during CFC (Mann–Whitney U Statistic: $^{+}P < 0.1$). **D.** Freezing of mice conditioned and re-exposed to the same context (A-A) and mice conditioned and exposed to a different context (A-B) during the 10 minutes test (two-way ANOVA: ns $P_{condition} > 0.05$). Aged mice display more generalisation. **E.** Schematic of contextual discrimination test. **F.** Freezing response during fear training. **G-H.** Freezing response (top) and discrimination index (bottom) at day 7 (G) and 8 (H) (upper graph: Mann–Whitney U Statistic; ns: non significant, $^{*}P < 0.05$, $^{**}P < 0.01$. Bottom graph: One-Sample Wilcoxon Test; ns: non significant, $^{*}P < 0.05$, $^{**}P < 0.01$).

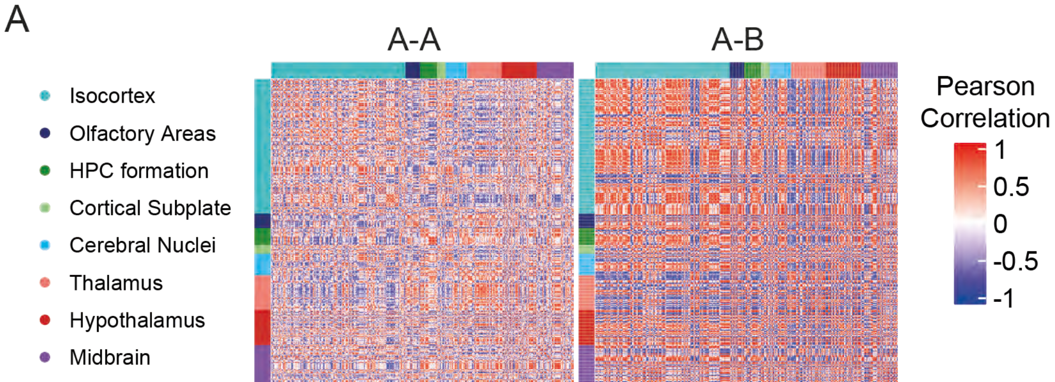

A
A-A
A-B
Isocortex
Olfactory Areas
HPC formation
Cortical Subplate
Cerebral Nuclei
Thalamus
Hypothalamus
Midbrain
Pearson Correlation
1
0.5
0
-0.5
-1

**Supplementary Figure S7 (related to Figure 3). Different patterns of engram activation in aged mice. A.** Subregion-by-subregion correlation matrices of brain activity, derived from pairwise comparisons of regional FOS expression patterns in each experimental group. Activity in each subregion is normalized to its volume.

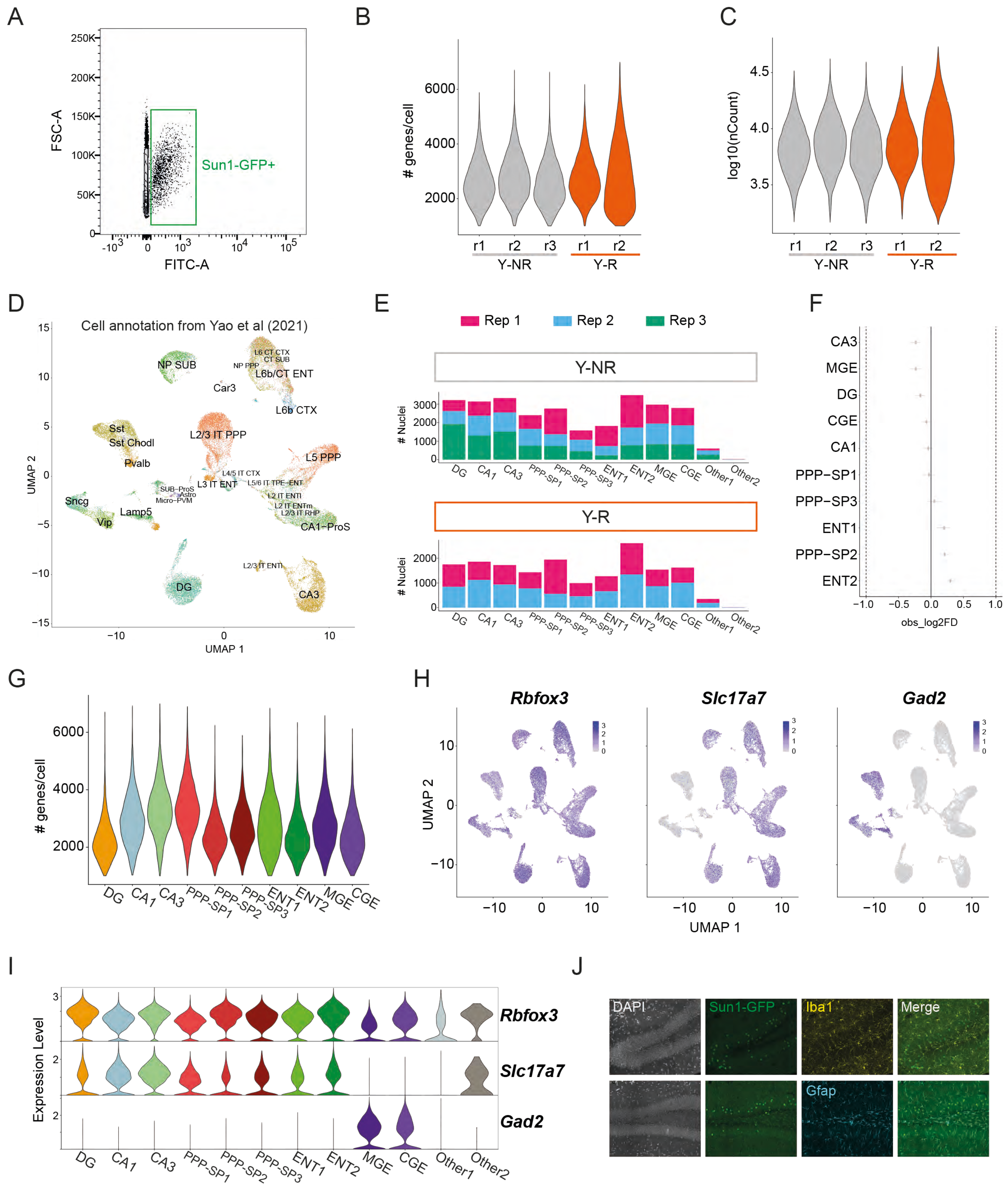
A
FSC-A
FITC-A
Sun1-GFP+
B
### genes/cell
r1 r2 r3 r1 r2
Y-NR
Y-R
C
log10(nCount)
D
Cell annotation from Yao et al (2021)
UMAP 1
UMAP 2
NP SUB
L6 CT CTX
CT SUB
NP PPP
L6b/CT ENT
Car3
L6b CTX
Sst
Sst Chodl
Pvalb
L2/3 IT PPP
L5 PPP
L4/5 IT CTX
L3 IT ENT
L5/6 IT TPE-ENT
SUB-ProS
Astro
Micro-PVM
L2 IT ENTl
L2 IT ENTm
L2/3 IT RHP
Sncg
Lamp5
Vip
CA1-ProS
L2/3 IT ENTl
DG
CA3
E
Rep 1
Rep 2
Rep 3
Y-NR
Y-R
### Nuclei
DG CA1 CA3 PPP-SP1 PPP-SP2 PPP-SP3 ENT1 ENT2 MGE CGE Other1 Other2
F
CA3
MGE
DG
CGE
CA1
PPP−SP1
PPP−SP3
ENT1
PPP−SP2
ENT2
obs_log2FD
G
### genes/cell
H
Rbfox3
Slc17a7
Gad2
UMAP 1
UMAP 2
I
Expression Level
Rbfox3
Slc17a7
Gad2
J
DAPI
Sun1-GFP
Iba1
Merge
Gfap

**Supplementary Figure S8 (related to Figure 4). Neuronal subtypes within the engram. A.** Flow cytometry analysis based on fluorescent intensity to distinguish cell populations by the presence or absence of SUN1-GFP. **B–C.** Violin plots showing the number of genes (**B**) and number of reads (**C**) per nucleus across biological replicates (rep1–rep3). **D.** UMAP visualization of cell type annotations based on the Yao et al. (2021) reference dataset[21]. **E.** Number of nuclei assigned to defined neuronal subtypes in each biological replicate for non-reactivated (NR, top) and reactivated (R, bottom) conditions. **F.** Permutation test assessing subtype enrichment. **G.** Number of genes detected per nucleus across neuronal subtypes. **H–I.** UMAP and violin plots showing expression of canonical markers: Rbfox3 (pan-neuronal), Slc17a7 (excitatory), and Gad2 (inhibitory); UMAP coloring reflects normalized expression levels. **J.** Microglial (Iba1) and astrocytic (Gfap) markers in the dentate gyrus do not colocalize with SUN1-GFP$^{+}$ cells.

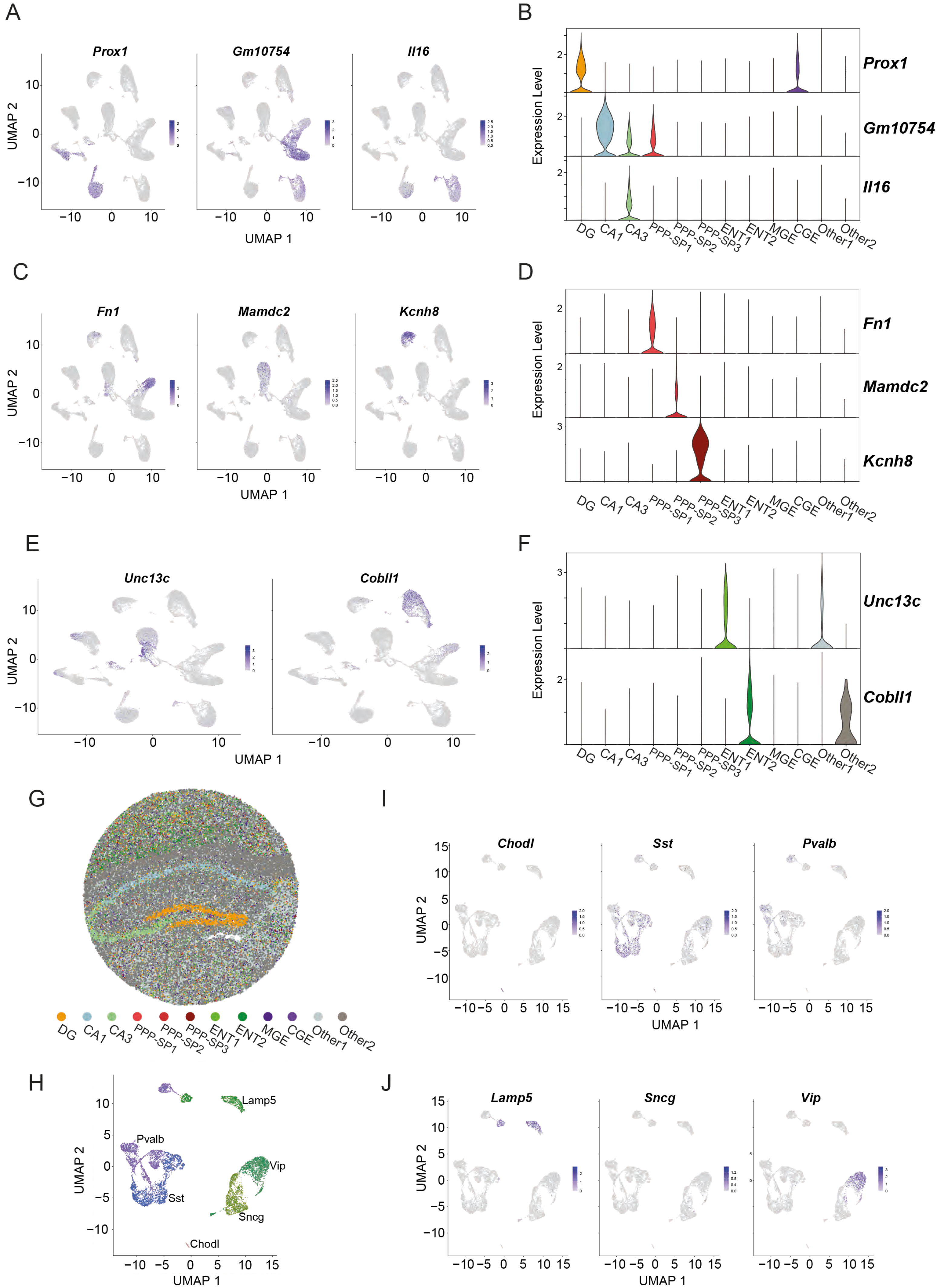
A
Prox1
Gm10754
Il16
UMAP 2
UMAP 1
B
Expression Level
Prox1
Gm10754
Il16
DG CA1 CA3 PPP-SP1 PPP-SP2 PPP-SP3 ENT1 ENT2 MGE CGE Other1 Other2
C
Fn1
Mamdc2
Kcnh8
UMAP 2
UMAP 1
D
Expression Level
Fn1
Mamdc2
Kcnh8
DG CA1 CA3 PPP-SP1 PPP-SP2 PPP-SP3 ENT1 ENT2 MGE CGE Other1 Other2
E
Unc13c
Cobll1
UMAP 2
UMAP 1
F
Expression Level
Unc13c
Cobll1
DG CA1 CA3 PPP-SP1 PPP-SP2 PPP-SP3 ENT1 ENT2 MGE CGE Other1 Other2
G
DG CA1 CA3 PPP-SP1 PPP-SP2 PPP-SP3 ENT1 ENT2 MGE CGE Other1 Other2
H
Lamp5
Pvalb
Vip
Sst
Sncg
Chodl
UMAP 2
UMAP 1
I
Chodl
Sst
Pvalb
UMAP 2
UMAP 1
J
Lamp5
Sncg
Vip
UMAP 2
UMAP 1

**Supplementary Figure S9 (related to Figure 4). Neuronal subtype markers within the engram. A–B.** UMAP and violin plots showing expression of excitatory neuron markers in the hippocampal region. **C–D.** UMAP and violin plots for markers of the PPP-SP class of excitatory neurons. **E–F.** UMAP and violin plots showing markers of excitatory neurons in the entorhinal cortex. **G.** Predicted spatial localization of cell types from our snRNA-seq data using RCTD mapping onto Slide-seqV2 hippocampus data[24]. **H.** UMAP displaying interneuron subtypes contributing to the engram. **I–J.** UMAP plots of interneuron populations, colored by normalized expression levels of specific subtype markers.

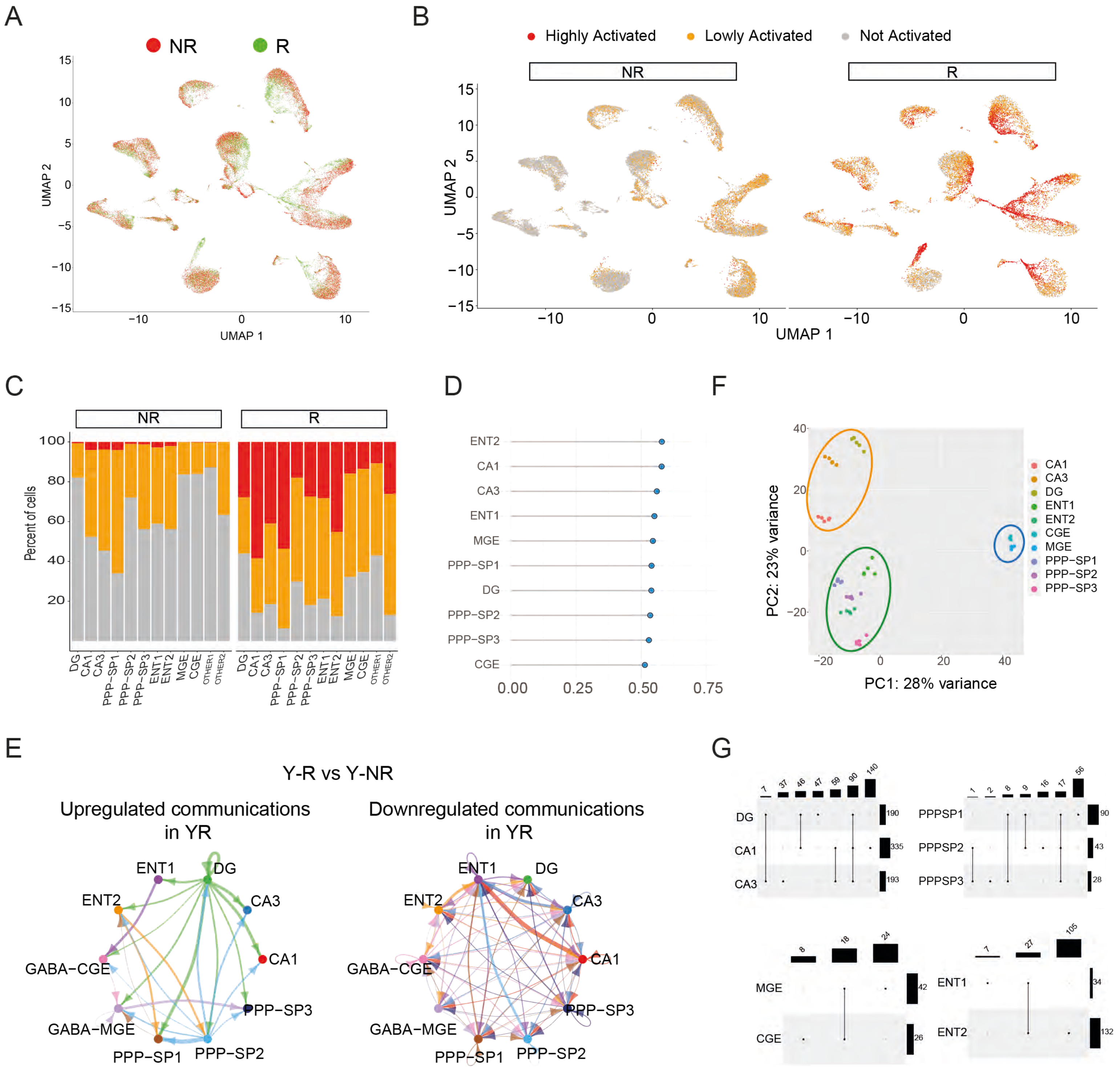
A
NR
R
UMAP 1
UMAP 2
B
Highly Activated
Lowly Activated
Not Activated
NR
R
UMAP 1
UMAP 2
C
NR
R
Percent of cells
DG
CA1
CA3
PPP-SP1
PPP-SP2
PPP-SP3
ENT1
ENT2
MGE
CGE
OTHER1
OTHER2
D
ENT2
CA1
CA3
ENT1
MGE
PPP-SP1
DG
PPP-SP2
PPP-SP3
CGE
F
PC1: 28% variance
PC2: 23% variance
CA1
CA3
DG
ENT1
ENT2
CGE
MGE
PPP-SP1
PPP-SP2
PPP-SP3
E
Y-R vs Y-NR
Upregulated communications in YR
Downregulated communications in YR
ENT1
DG
ENT2
CA3
CA1
GABA-CGE
GABA-MGE
PPP-SP3
PPP-SP1
PPP-SP2
G
DG
CA1
CA3
PPPSP1
PPPSP2
PPPSP3
MGE
CGE
ENT1
ENT2

**Supplementary Figure S10 (related to Figure 4). Memory retrieval elicits distinct transcriptional responses across neuronal subtypes. A.** UMAP plot of SUN1-GFP$^+$ nuclei in the R (green) and NR (red) groups based on snRNA-seq data. **B.** UMAP plots for NR (left) and R (right) groups showing the inferred activation state of each nucleus. **C.** Proportions of activated nuclei within each neuronal subtype for NR and R groups. **D.** *Augur* analysis: area under the receiver operating characteristic curve (AUC) values exceed 0.5 for all cell types, indicating discriminatory transcriptional responses. **E.** Circle plots generated by NeuronChat depicting upregulated (left) and downregulated (right) intercellular communication in the Y-R versus Y-NR comparison. **F.** PCA of pseudobulk transcriptomes from each replicate across neuronal subtypes. **G.** UpSet plot illustrating intersections among DEGs across neuronal types.

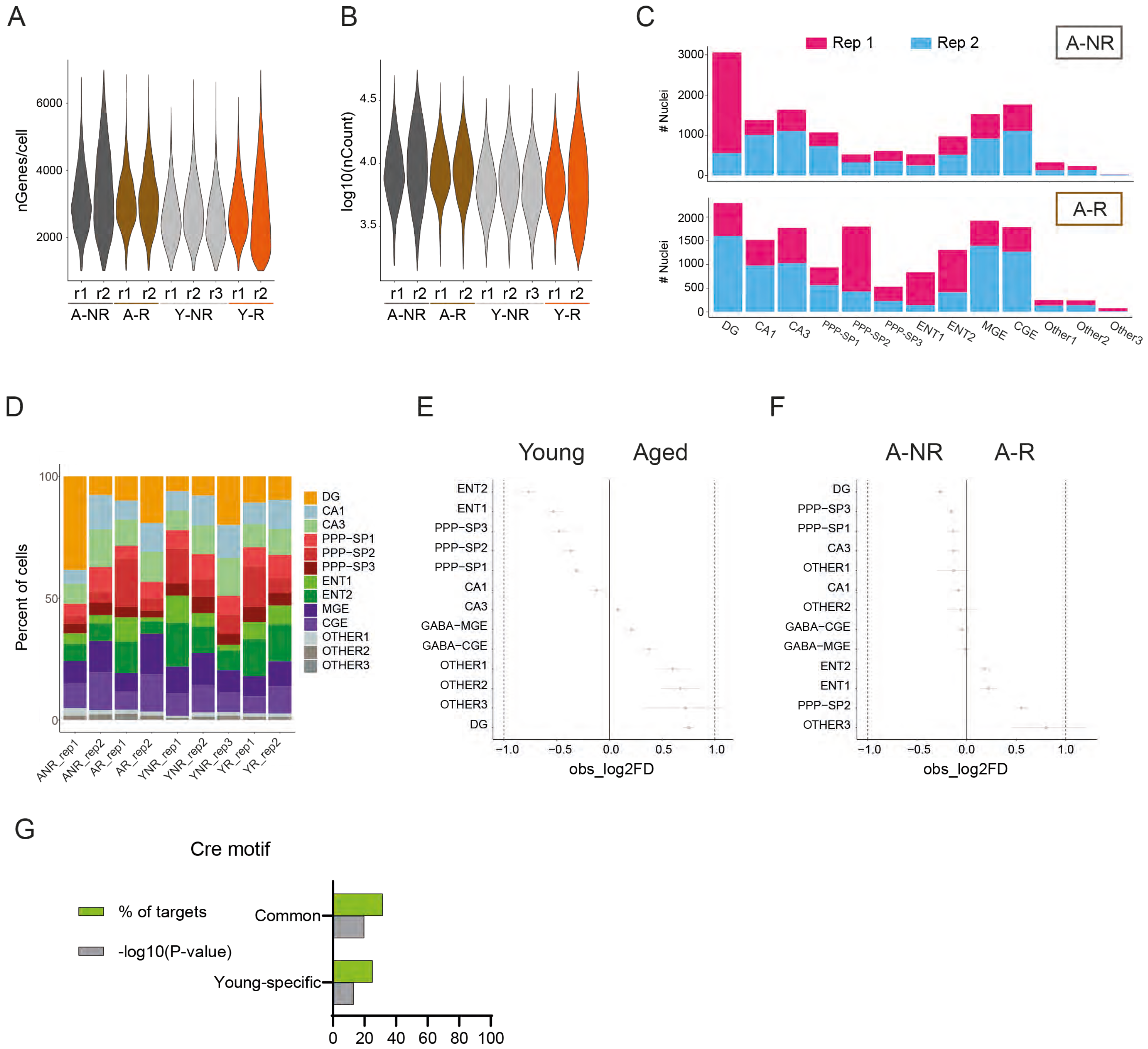

A
nGenes/cell
6000
4000
2000
r1 r2 r1 r2 r1 r2 r3 r1 r2
A-NR A-R Y-NR Y-R
B
log10(nCount)
4.5
4.0
3.5
r1 r2 r1 r2 r1 r2 r3 r1 r2
A-NR A-R Y-NR Y-R
C
Rep 1
Rep 2
A-NR
A-R
### Nuclei
DG CA1 CA3 PPP-SP1 PPP-SP2 PPP-SP3 ENT1 ENT2 MGE CGE Other1 Other2 Other3
D
Percent of cells
DG
CA1
CA3
PPP−SP1
PPP−SP2
PPP−SP3
ENT1
ENT2
MGE
CGE
OTHER1
OTHER2
OTHER3
ANR_rep1 ANR_rep2 AR_rep1 AR_rep2 YNR_rep1 YNR_rep2 YNR_rep3 YR_rep1 YR_rep2
E
Young Aged
ENT2
ENT1
PPP−SP3
PPP−SP2
PPP−SP1
CA1
CA3
GABA−MGE
GABA−CGE
OTHER1
OTHER2
OTHER3
DG
obs_log2FD
F
A-NR A-R
DG
PPP−SP3
PPP−SP1
CA3
OTHER1
CA1
OTHER2
GABA−CGE
GABA−MGE
ENT2
ENT1
PPP−SP2
OTHER3
obs_log2FD
G
Cre motif
% of targets
-log10(P-value)
Common
Young-specific
0 20 40 60 80 100

**Supplementary Figure S11 (related to Figure 5). Neuronal composition of engram cells in young and aged mice. A–B.** Violin plots showing the number of genes (**A**) and number of reads (**B**) per nucleus across biological replicates (rep1–rep3). **C.** Number of nuclei from each biological replicate in aged mice annotated as defined neuronal subtypes for NR (top) and R (bottom) conditions. **D.** Percentage of nuclei assigned to each neuronal subtype in young and aged mice, across biological replicates. **E.** Permutation test comparing neuronal subtype distributions between young and aged mice. **F.** Permutation test comparing R and NR conditions within aged mice. **G.** Transcription factor motif analysis.

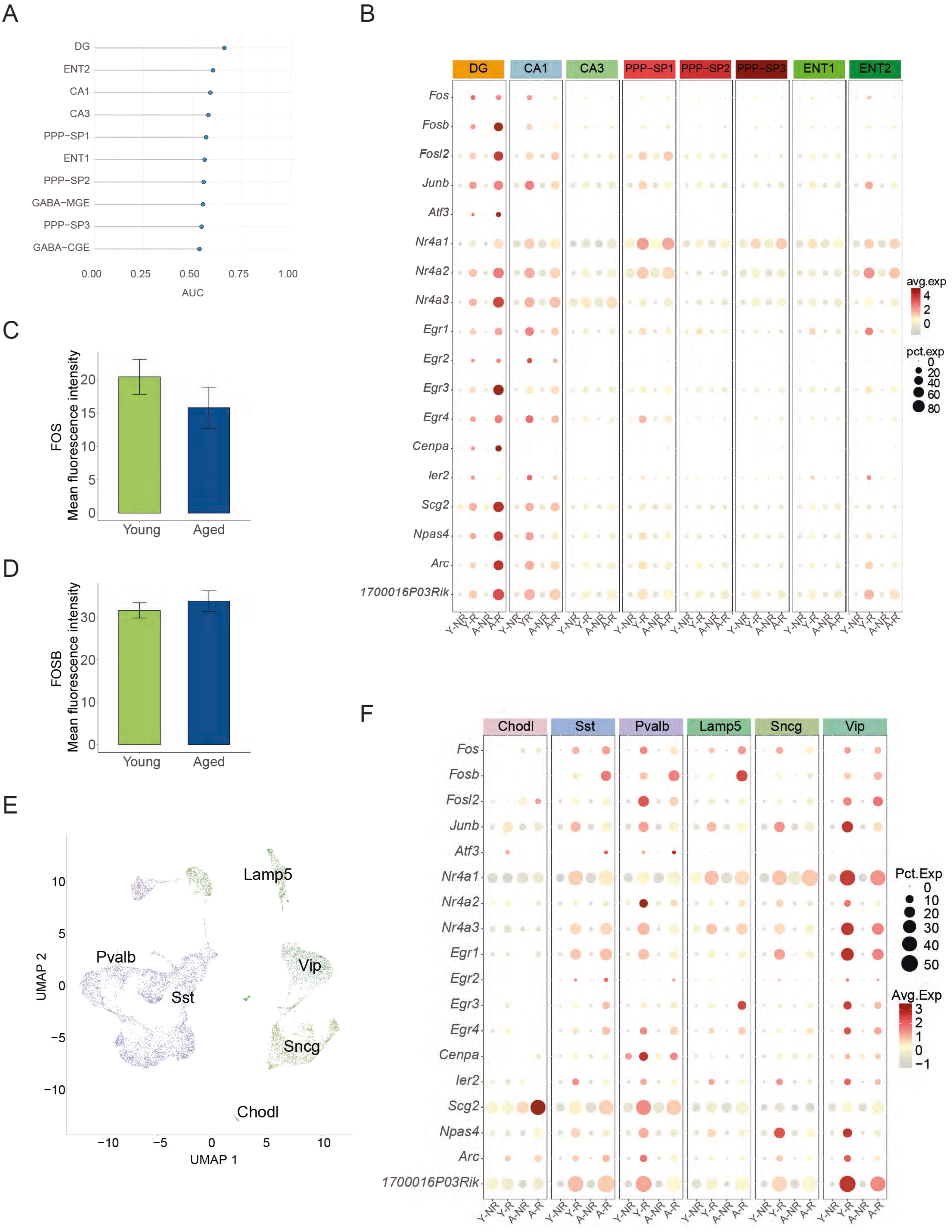
A
DG
ENT2
CA1
CA3
PPP−SP1
ENT1
PPP−SP2
GABA−MGE
PPP−SP3
GABA−CGE
0.00
0.25
0.50
0.75
1.00
AUC
B
DG
CA1
CA3
PPP−SP1
PPP−SP2
PPP−SP3
ENT1
ENT2
Fos
Fosb
Fosl2
Junb
Atf3
Nr4a1
Nr4a2
Nr4a3
Egr1
Egr2
Egr3
Egr4
Cenpa
Ier2
Scg2
Npas4
Arc
1700016P03Rik
Y-NR
Y-R
A-NR
A-R
avg.exp
4
2
0
pct.exp
0
20
40
60
80
C
FOS
Mean fluorescence intensity
0
5
10
15
20
Young
Aged
D
FOSB
Mean fluorescence intensity
0
10
20
30
Young
Aged
E
Lamp5
Pvalb
Sst
Vip
Sncg
Chodl
UMAP 2
UMAP 1
10
5
0
−5
−10
F
Chodl
Sst
Pvalb
Lamp5
Sncg
Vip
Pct.Exp
0
10
20
30
40
50
Avg.Exp
3
2
1
0
−1

**Supplementary Figure S12 (related to Figure 5). Transcriptional responses to memory recall in young and aged mice. A.** *Augur* analysis showing that the area under the receiver operating characteristic curve (AUC) exceeds random chance (0.5) for all cell types. **B.** Bubble plot of key IEGs across excitatory neuron subtypes. Color indicates average normalized expression; circle size reflects the percentage of nuclei expressing the gene across conditions: Y-NR (young, no recall), Y-R (young, recall), A-NR (aged, no recall), A-R (aged, recall). **C–D.** Comparison of mean fluorescence intensity of FOS (**C**) and FOSB (**D**) in SUN1-GFP$^+$FOS$^+$ nuclei between young and aged mice. Bars represent mean ± SEM; significance was determined using independent samples *t*-tests. **E.** UMAP plot of inhibitory interneuron subtypes. **F.** Bubble plot of key IEGs across inhibitory neuron subtypes, with color and circle size representing normalized expression and expression frequency, respectively, for each experimental condition (Y-NR, Y-R, A-NR, A-R).

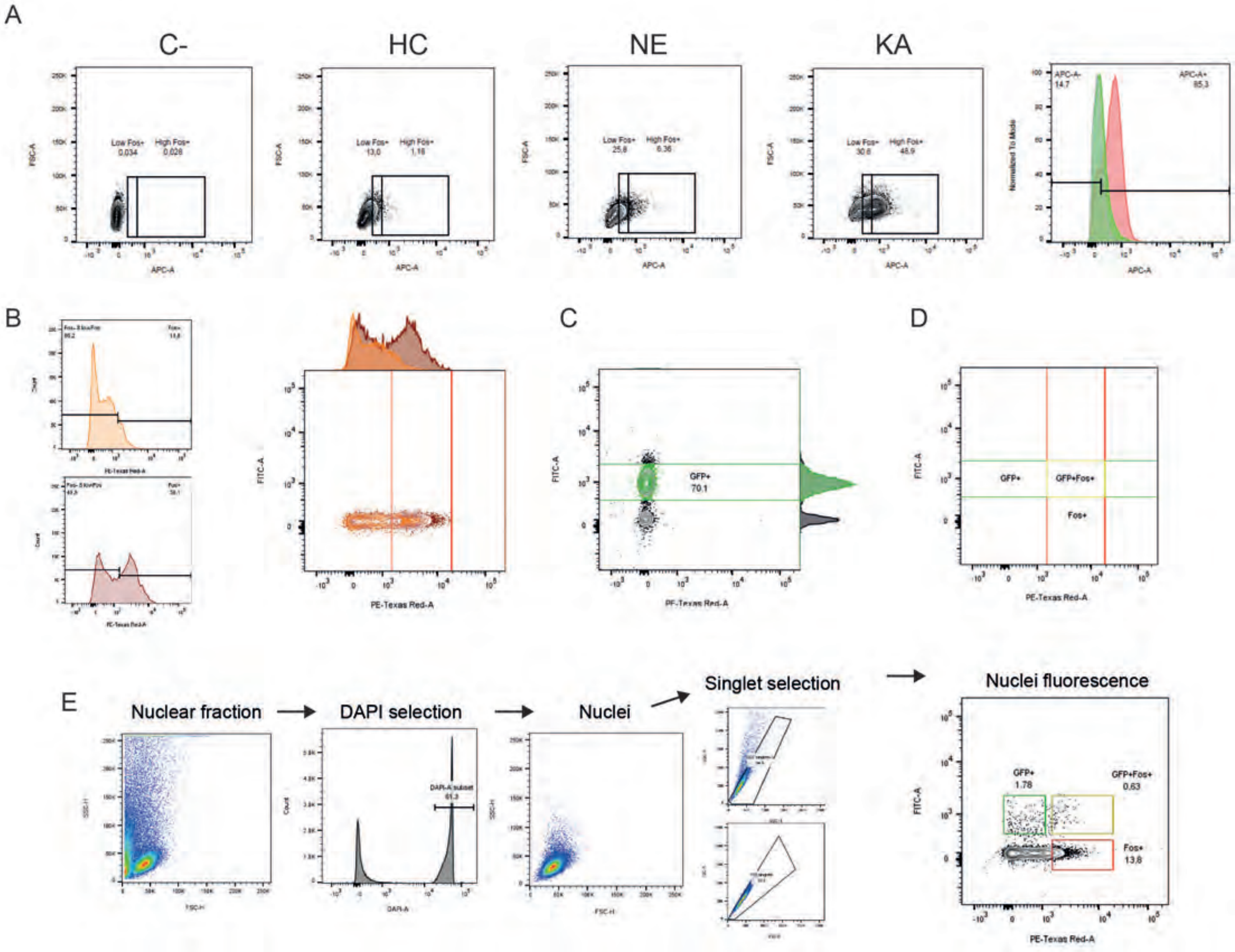
A
C-
HC
NE
KA
Low Fos+
High Fos+
0,034
0,028
13,0
1,16
25,8
6,36
30,6
48,9
APC-A-
14,7
APC-A+
85,3
FSC-A
APC-A
Normalized To Mode
B
C
D
FITC-A
PE-Texas Red-A
GFP+
70,1
GFP+Fos+
Fos+
E
Nuclear fraction
DAPI selection
Nuclei
Singlet selection
Nuclei fluorescence
DAPI-A subset
61,3
SSC-H
FSC-H
DAPI-A
Count
1.78
0.63
13,8

**Supplementary Figure S13 (related to Figure 6). Flow cytometry–based isolation of engram cells. A.** Representative FACS plots showing FOS expression (x-axis) across nuclear populations following home cage (HC), novel environment exposure (NE), or kainate (KA) administration. Percentages of low- and high-FOS–expressing nuclei are indicated above the gates. **B.** Percentage of $FOS^{+}$ nuclei following fear memory recall (orange) and KA-induced activation (red). **C.** Percentage of SUN1-$GFP^{+}$ nuclei in CaMKII-CreERT2 mice crossed with a Cre-dependent SUN1-GFP reporter line after TMX administration. **D.** Gating strategy for sorting three defined nuclear populations. **E.** Stepwise gating and channel-based filtering used to isolate fluorescent singlet nuclei by flow cytometry. Percentages indicate the proportion of events retained at each gating step.

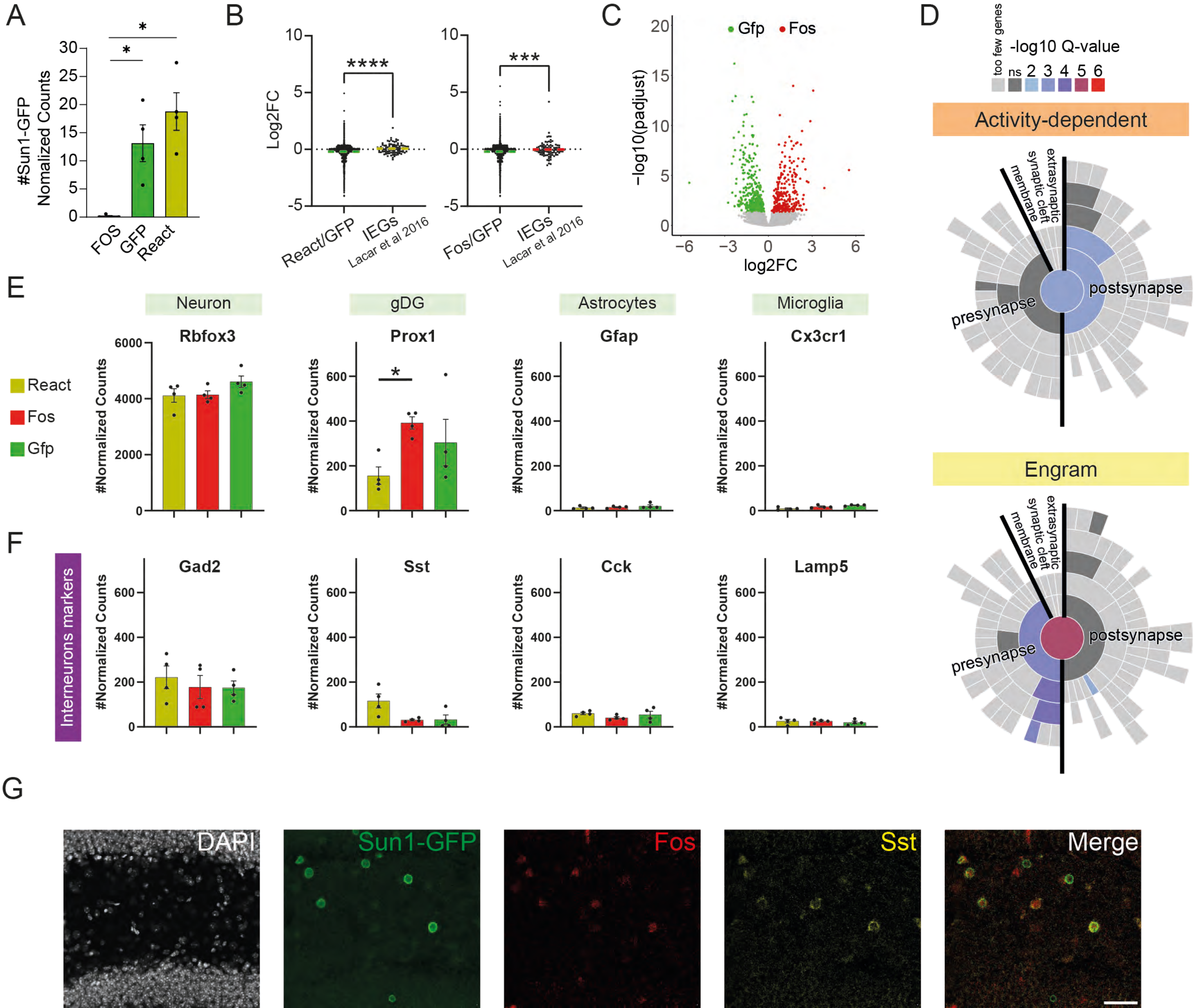
A
#Sun1-GFP Nomalized Counts
FOS
GFP
React
B
Log2FC
React/GFP
IEGs Lacar et al 2016
Fos/GFP
IEGs Lacar et al 2016
C
Gfp
Fos
−log10(padjust)
log2FC
D
too few genes
-log10 Q-value
ns 2 3 4 5 6
Activity-dependent
extrasynaptic
synaptic cleft
membrane
presynapse
postsynapse
Engram
E
Neuron
gDG
Astrocytes
Microglia
Rbfox3
Prox1
Gfap
Cx3cr1
React
Fos
Gfp
#Normalized Counts
F
Interneurons markers
Gad2
Sst
Cck
Lamp5
G
DAPI
Sun1-GFP
Fos
Sst
Merge

**Supplementary Figure S14 (related to Figure 6). Transcriptional programs and cellular composition of reactivated cells in the DG. A.** Normalized counts of SUN1-GFP expression (Mann–Whitney *U* test). **B.** Log2FC of all genes (green) in the comparisons of Reactivated vs 1st Experience (left) and 2nd Experience vs 1st Experience (right). Highlighted in yellow and red are genes most strongly induced in $FOS^+$ vs $FOS^-$ nuclei, based on Lacar et al.[58]. **C.** Volcano plot showing significance ($-\log_{10}$ *p*-adjusted) and log2FC distribution for differentially expressed genes between 1st and 2nd Experience cells. **D.** Enrichment of synaptic Gene Ontology (GO) terms (SynGO Cellular Component) among genes from activity-dependent and engram-specific clusters. **E.** Normalized counts of canonical cell type markers (Mann–Whitney *U* test). **F.** Normalized counts of interneuron subtype markers. **G.** Representative confocal images of reactivated somatostatin-positive interneurons in the hilus. Scale bar: 50 μm. $^*P < 0.05$, $^{**} P < 0.01$, $^{***} P < 0.001$.

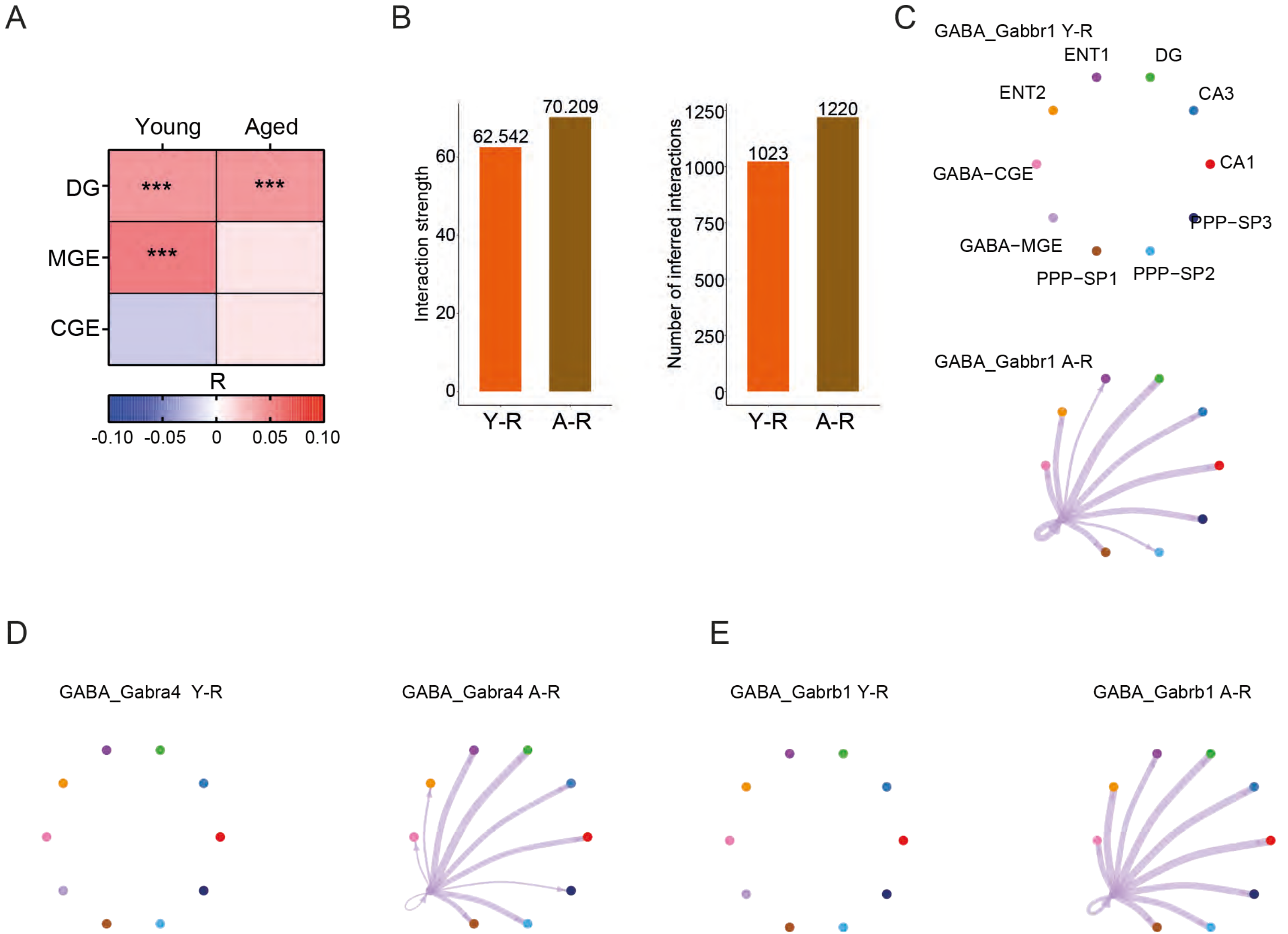

A
Young
Aged
DG
MGE
CGE
***
***
***
R
-0.10
-0.05
0
0.05
0.10
B
Interaction strength
62.542
70.209
Y-R
A-R
Number of inferred interactions
1023
1220
C
GABA_Gabbr1 Y-R
ENT1
DG
ENT2
CA3
GABA−CGE
CA1
PPP−SP3
GABA−MGE
PPP−SP1
PPP−SP2
GABA_Gabbr1 A-R
D
GABA_Gabra4 Y-R
GABA_Gabra4 A-R
E
GABA_Gabrb1 Y-R
GABA_Gabrb1 A-R

**Supplementary Figure S15 (related to Figure 7). Increased GABAergic connectivity in aged mice. A.** Heatmap showing Pearson's correlation between snRNA-seq data from DG, GABA-MGE, and GABA-CGE clusters (log2FC activated vs non activated) and nuRNA-seq data (log2FC reactivated vs 1$^{st}$ experience) in young and aged mice. **B.** Comparison of the strength (left) and number (right) of inferred cell–cell interactions during memory recall in young and aged mice. **C–E.** Network diagrams depicting individual inhibitory communication via GABA receptors: GABBR1 (**C**), GABRA4 (**D**), and GABRB1 (**E**).
